# Live cell imaging and analysis reveal cell phenotypic transition dynamics inherently missing in snapshot data

**DOI:** 10.1101/2019.12.12.874248

**Authors:** Weikang Wang, Diana Douglas, Jingyu Zhang, Yi-Jiun Chen, Ya-Yun Cheng, Sangeeta Kumari, Metewo Selase Enuameh, Yan Dai, Callen T. Wallace, Simon C. Watkins, Weiguo Shu, Jianhua Xing

## Abstract

Recent advances in single-cell techniques catalyze an emerging field of studying how cells convert from one phenotype to another, in a step-by-step process. Two grand technical challenges, however, impede further development of the field. Fixed cell-based approaches can provide genome-wide snapshots of cell status but have fundamental limits on revealing temporal information, and fluorescence-based live cell imaging approaches provide temporal information but are technically challenging for multiplex long-term imaging. We first developed a live-cell imaging platform that tracks cellular status change through combining endogenous fluorescent labeling that minimizes perturbation to cell physiology, and/or live cell imaging of high-dimensional cell morphological and texture features. With our platform and an A549 VIM-RFP EMT reporter line, live cell trajectories reveal parallel paths of epithelial-to-mesenchymal transition missing from snapshot data due to cell-cell heterogeneity. Our results emphasize the necessity of extracting dynamical information of phenotypic transitions from multiplex live cell imaging.

## Introduction

Cells of a multicellular organism can assume different phenotypes that can have drastically different morphological and gene expression patterns. A fundamental question in developmental biology is how a single fertilized egg develops into different cell types in a spatial-temporally controlled manner. Cell phenotypic transition (CPT) also takes place for differentiated cells under physiological and pathological conditions. A well-studied example is the epithelial-to-mesenchymal transition (EMT), central to many fundamental biological processes including embryonic development and tissue regeneration, wound healing, and disease-like states such as fibrosis and tumor invasiveness (*1*). One additional example is artificially reprogramming differentiated cells, such as fibroblasts, into induced pluripotent stem cells and other differentiated cell types such as neurons and cardiomyocytes (*2*). CPT is ubiquitous in biology, and a mechanistic understanding of how a CPT proceeds, emerges as a focused research area with an ultimate goal of achieving effective control of the phenotype of a cell.

The CPT studies can also be placed in another large context of studying dynamical processes of escaping from a metastable state or relaxing to a newly-established stationary state (*3*). Such problem has been a focused and still active research topic in physics and chemistry for more than a century. Formulating CPT as such a problem allows one to apply techniques such as modern transition path sampling and transition path theory (*4*), and control theories of dynamical systems, for mechanistic understanding of the transition bottlenecks, and manipulating the transitions such as accelerating and directing reprogramming and differentiation processes, and slowing down/preventing undesirable pathological processes. In return CPT studies can provide unprecedented multi-dimensional information that is difficult to obtain for a molecular system, and thus ideal for testing and advancing theoretical studies of non-equilibrium transition processes. A framework of quantitative description of CPT processes is necessary to catalyze such epochal convergence of traditionally separated fields.

Recent advances in snapshot single cell techniques, notably single cell RNA-seq and imaging-based techniques, pose potential questions, such as how does a CPT process proceed, step-by-step, along the continuous high-dimensional gene expression (e.g., transcriptome, proteome) space. These destructive methods, however, are inherently unable to reveal the temporal dynamics of how an individual cell evolves over time during a CPT. Approaches such as pseudo-time trajectory analysis (*5*), ergodic rate analysis (*6*) and RNA velocities (*7*) have been developed to retrieve partial dynamical information from snapshot data.

However, some fundamental limits exist in inferring dynamical information from snapshot data (*8*). Figure 1 illustrates that inference from snapshot data unavoidably misses key dynamical features. A bi-stable system is coupled to a hidden process, e.g., epigenetic modification for a cellular system, which is slow compared to the transition process being studied (Fig. 1A). Presence of the slow variable leads to observed heterogeneous dynamics (Fig. 1B, more details of the simulation are in Fig. S1). Individual trajectories show characteristic stepping dynamics, but the transition positions vary among different trajectories due to the system assuming different values for the hidden slow variable. Consequently, when only snapshot data is available, information about the temporal correlation of individual cell trajectories is missing, and one cannot deduce the stepping dynamics (Fig. 1C).

**Fig. 1.**
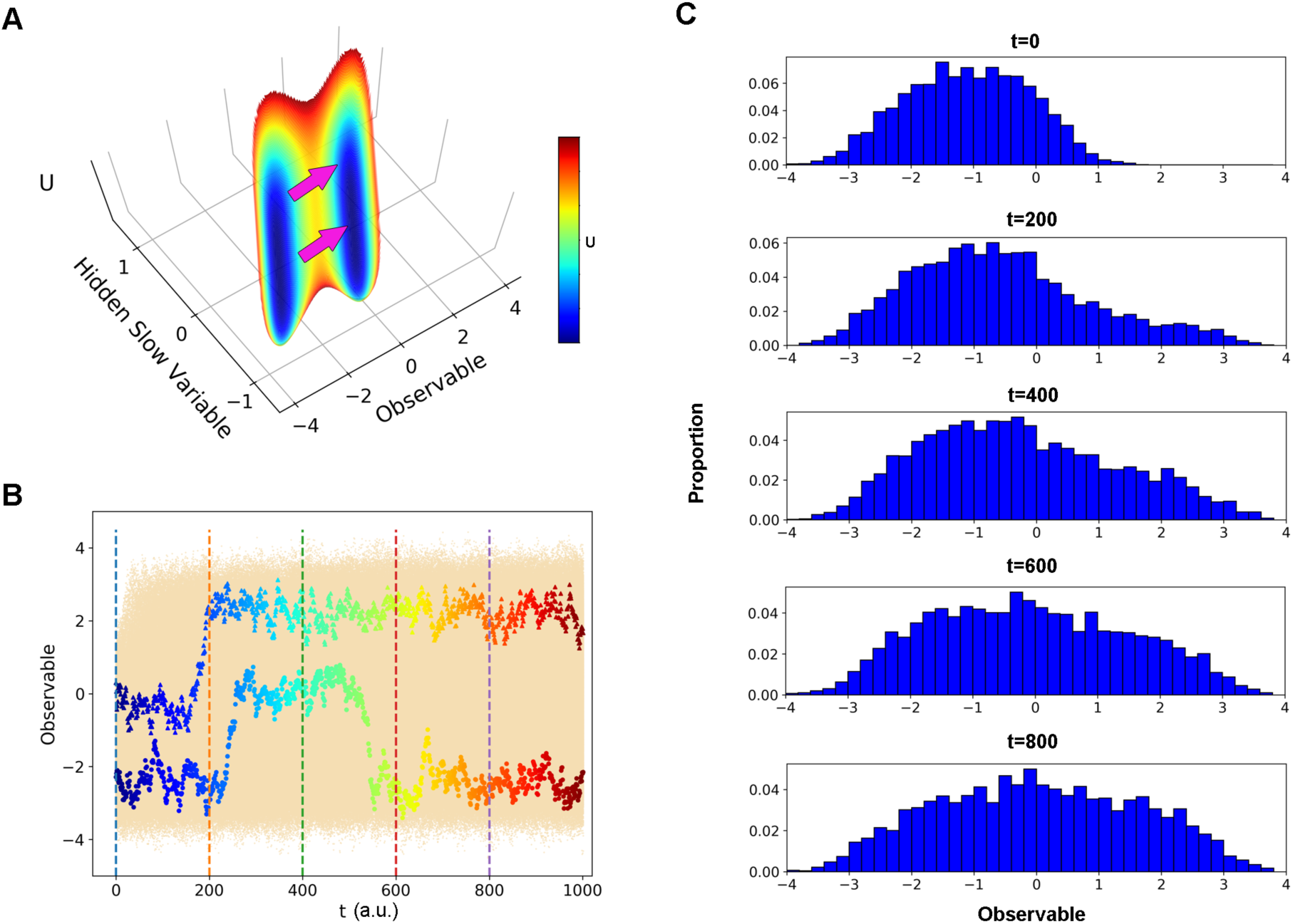
Hidden slow variables can conceal some dynamical features from snapshot data. (**A**) A double well potential with one observable coupled to a hidden variable with dynamics much slower than that of the process under study. **(B)** Superimposed trajectories from stochastic simulation, with two typical trajectories highlighted under different values of hidden slow variable. Color represents time. **(C)** Histogram of the observable at various time points, reflecting the snapshot data. The snapshot data does not follow a bimodal distribution characteristic of the stepping dynamics due to cell-cell heterogeneity from the coupling between hidden slow variable and observable.

The necessity of live cell trajectories has been illustrated in a number of studies, such as the information capacity of a signal transduction network (*9*), incoherent-feedforward loops to detect only fold change but not the absolute change of an input signal (*10*), and step-wise cellular responses to drugs (*11*). Two recent studies conclude a linear path for EMT from analyzing single cell RNA-seq and proteomic data (*12, 13*). This conclusion, however, is inconsistent with theoretical predictions of parallel paths for a multi-stable system like EMT (*14, 15*), raising a question of whether the linear path is an artifact from snapshot data. Live cell imaging is needed to address such question.

Therefore, acquiring information from long-term CPT dynamics requires tracking individual cells through live cell imaging, typically with time-lapse fluorescent imaging. However, identifying appropriate species that faithfully reflect the process for labeling, and generating such labeling, can be tedious and time-consuming. Additionally, multiplex and frequent fluorescent image acquisition over a long period of time, e.g., days, is necessary for characterizing a CPT process, but is severely limited by the number of available fluorescence channels and cytotoxicity concerns.

In short, a technical dilemma exists: fixed cell-based techniques provide high-dimensional expression profiles of individual cells but lack true dynamical information, while fluorescent-labeling based live cell imaging techniques typically provide dynamical information for only a small number of dynamical variables. To tackle the substantial challenges in CPT studies, here we develop a framework for extracting cell dynamical information through quantitative analysis of live cell trajectories in a combined high dimensional cell morphology and expression space. Our method resides on the observation that either the expression pattern or cell morphological features can define a cell state. The latter broadly refers to collective cellular properties such as cell body shape, organelle distribution, etc., which are convenient for live cell imaging, with and without labeling. Hundreds of such morphological features have been routinely used in pathology, and in a number of fixed and live cell studies for defining and studying cell phenotypes, and drug responses (*11*). Introduction of this framework allows one to study CPTs in the context of well-established chemical reaction rate theories, specifically the transition path theory and transition path sampling as mentioned above (*4, 16*).

We applied this framework to study TGF-β induced EMT in a human A549 cell line with endogenous vimentin-RFP labeling. We represented cell states in a composite 309 dimensional feature space of the cell body contour shape and distribution of vimentin, an intermediate filament and a key mesenchymal marker. While the framework is for morphological features in general, in this study we focus on the cell body shape, and use cell shape and morphology indistinctively. Through quantifying time-lapse images, aided by a deep learning based image analysis algorithm, we were able to define the epithelial and mesenchymal states and unravel two parallel pathways that EMT proceeds through. We provide a Python package, Multiplex Trajectory Recording and Analysis of Cellular Kinetics, or M-TRACK, for studying CPT in the morphology/expression space. The framework will provide a foundation for quantitative experimental and theoretical studies of CPT dynamics.

## Results

### Human A549 cells with endogenous vimentin-RFP labeling were generated using the CRISPR-Cas9 system

The A549 VIM-RFP cell line was created using CRISPR/Cas9 technology, in which the red fluorescence protein (RFP) sequence was inserted just before the stop codon of the endogenous vimentin gene (Fig 2A). The VIM-RFP knock-in allele was confirmed by co-localization immuno-staining of vimentin and VIM-RFP (Fig. 2B), sequencing (Fig. S2A), and Western blot (Fig. S2B). With continuous treatment of recombinant human TGF-β1 for two days, most cells underwent apparent cell shape changes from round polygon shapes to elongated spear shapes. The A549 VIM-RFP cells have basal vimentin expression, suggesting the cells have already undergone partial EMT, as previously reported for the parental A549 cells (*13*). With TGF-β treatment, the A549 VIM-RFP cells show increased invasive capacity reflecting the functionality of mesenchymal cells (Fig. 2C). The immuno-staining results showed that Snail1 and N-cadherin increased their expression (Fig. S2C), further confirming the occurrence of EMT.

**Fig. 2.**
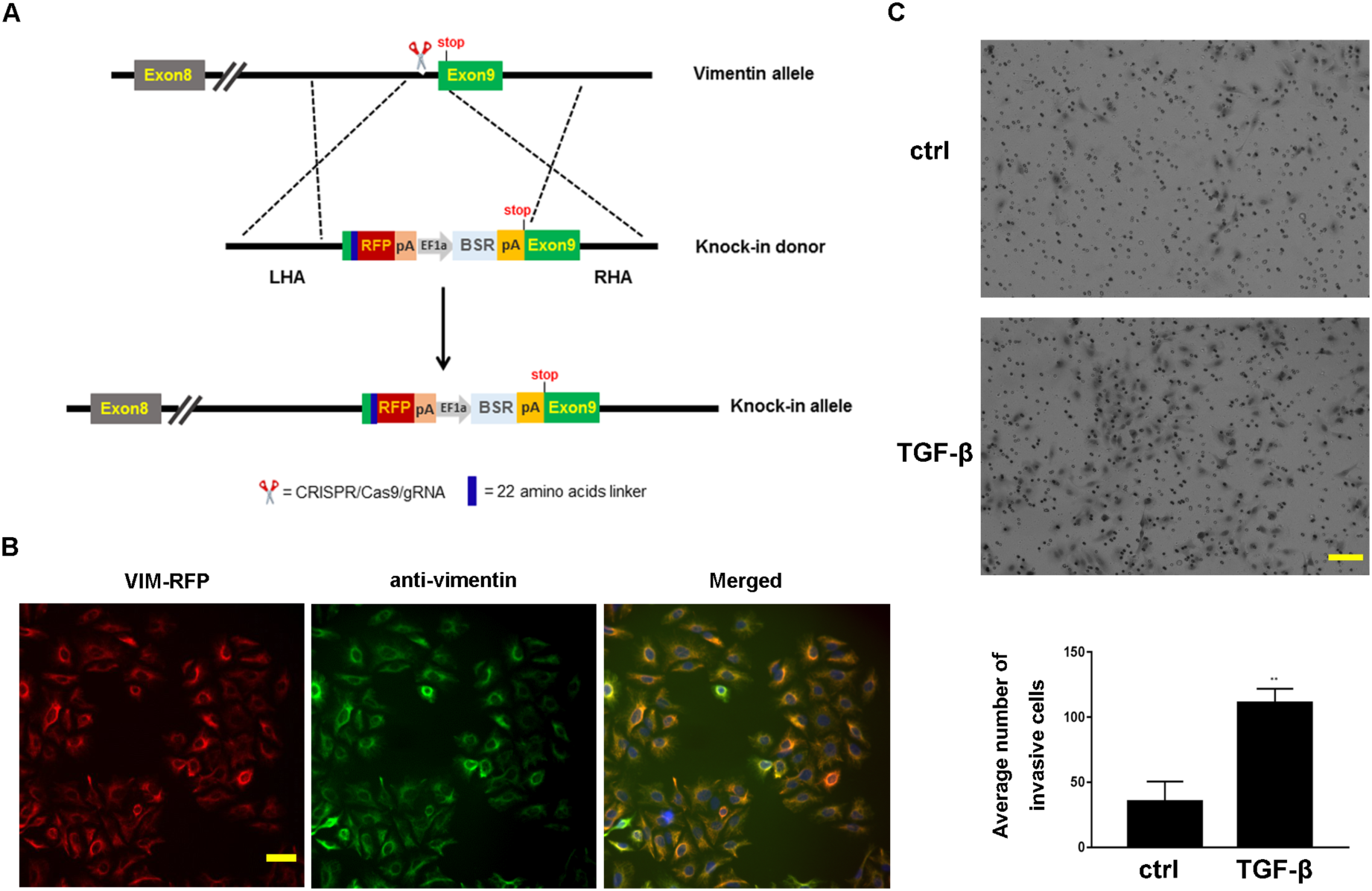
The generated A549 vimentin-RFP cell line was confirmed for physiology. (**A**) The Vimentin RFP knock-in donor. Schema of CRISPR-Cas9 mediated generation of VIM-RFP knock-in allele. (**B**) Colocalization immuno-staining of vimentin and endogenous vimentin RFP, scale bar equals 20 µm. (**C**) Matrigel invasion assay of A549 vimentin-RFP cells. After TGF-β induction, cells show increased invasive capacity, scale bar equals 100µm.

### A complete orthonormal basis set represents cell body contour shapes

To mathematically describe how a CPT (EMT here) proceeds, one needs to choose a mathematical representation of cell status at a given time. For representing the cell shape, we adopted the active shape model that has been widely used in computer based image analyses (*17*), and particularly in cell biology studies (*18, 19*), but here, we use it for the purpose of forming a complete orthonormal basis set (Fig. 3). That is, we first segmented the images using our modified deep convolutional neural network procedure (Fig. 3A, see Supplemental Text for details) and tracked individual cell trajectories. Each cell shape was aligned to a reference shape and was approximated by *N (= 150)* landmark points equally spaced along the cell contour (*20*) (Fig. 3B). For two-dimensional images, a cell was specified by a point **z** = (x_1_, x_2_, …, x_N_; y_1_, y_2_, …, y_N_) in the 2*N* dimensional shape space (Fig. 3C). By performing principal component analysis (PCA) on the data set of a collection of single cell trajectories, one constructs a complete orthonormal basis set with the 2*N* − 4 eigenvectors {*a*}, or the principal modes, for the shape space. Notice that alignment fixes four degrees of freedom (center and orientation). An attractive feature of the principal modes is that they have clear physical meanings. For example, the two leading principal modes of A549 VIM-RFP cells undergoing EMT reflect cell growth along the long and short axes, respectively (Fig. 3D). Then any cell shape that is approximated by the landmark point **z**’(t), which is generally time dependent in live cell imaging, can be expressed as a linear combination of these principal modes, 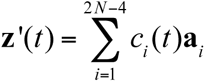 (Fig. 3E). Therefore, **c**(t) =(c_1_(t),…, c_2N-4_(t)) forms a trajectory in the shape space expanded by the basis set {*a*}.

**Fig. 3.**
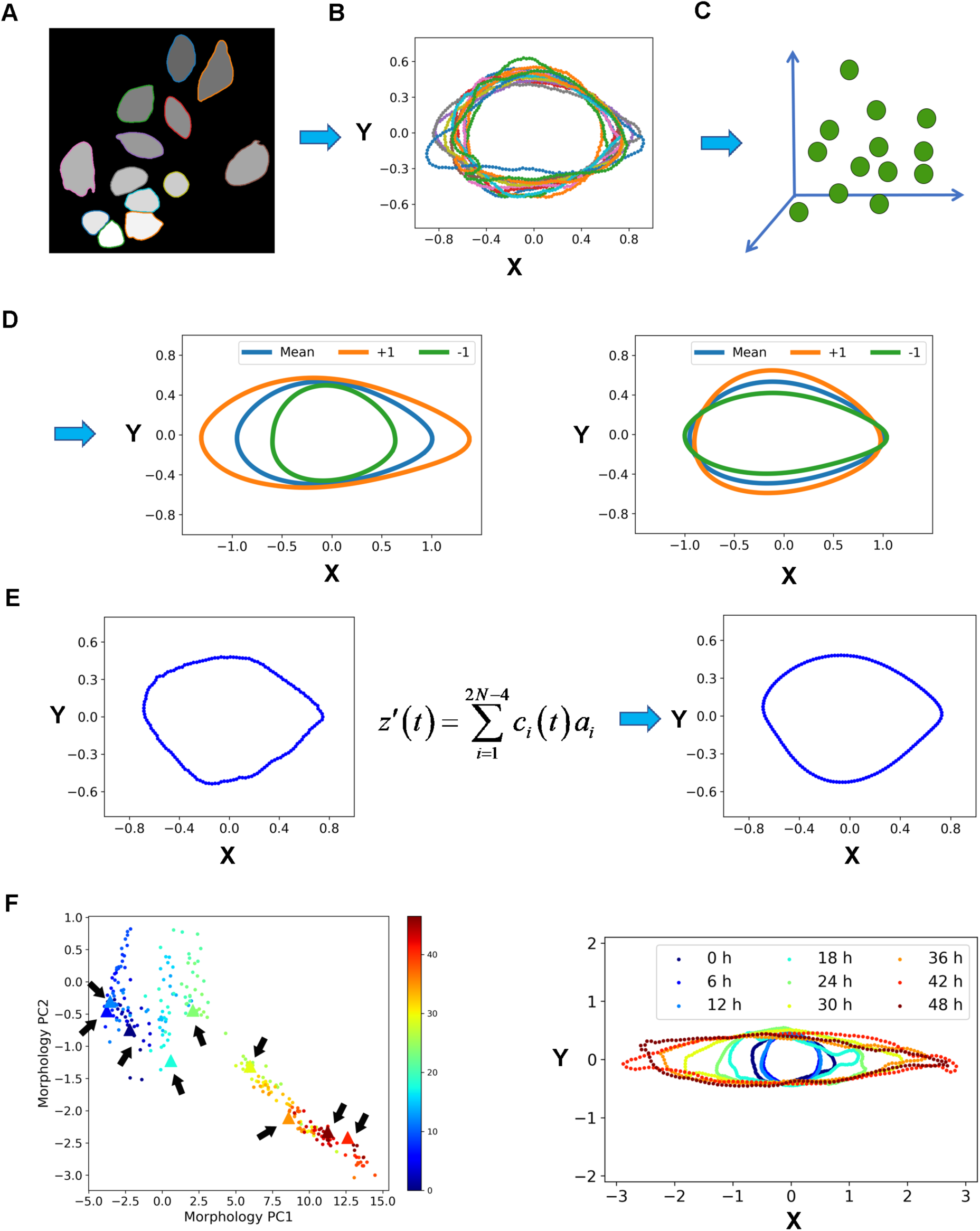
Single cell trajectories are quantitative represented in the morphology space. (**A**) Segmentation of single cells with the DCNN/watershed method. (**B**) Extraction of cell outline, alignment to a mean cell shape, and resampling using the active shape model. (**C**) Representation of single cell shapes as a point in the 2N - 4 dimensional morphology space. (**D**) Principal modes of variation of morphology. Left: First principal mode; Right: Second principal mode. +1 represents the corresponding morphology’s coordinate value on the axis of PC1 or PC2 is 1. -1 represents the corresponding morphology’s coordinate value on the axis of PC1 or PC2 is -1. The principal modes reflects the characteristics of cell morphology variation along the axis of PCs. (**E**) Reconstruction of cell shapes with principal modes. (**F**) A typical single cell trajectory in the leading morphology PC domain (left) and its corresponding contours (triangle dots marked by arrows in the left that have the same color as the contours) at various time points (right). Color bar represent time (unit in hour).

Figure 3F shows a typical trajectory projected to the first two leading principal component (PC) modes in the shape space and their corresponding time courses of cell contour shape changes. Over time this cell elongated along the major axis, (PC1) while shortened slightly along the minor axis, resulting in a long rod shape with enlarged cell size. Two additional trajectories in Fig. S3 further reveal that single cell trajectories are heterogeneous with switch-like or continuous transitions, while sharing similar elongation of PC1 over time.

### Haralick features quantify texture feature change of cytosolic distribution of vimentin during EMT

During a CPT, cell morphology changes are accompanied by global changes in gene expression profiles (*21*). Specifically, in A549 VIM-RFP cells that have been treated with TGF-β for two days, vimentin was upregulated with a texture structure change from being condensed in certain regions of the cell to be dispersed throughout the cytosol (Fig. 4A), consistent with previous reports (*22–24*). These previous studies used fixed cells, and lack temporal information about the vimentin dynamics. Therefore, we recorded the change in vimentin within individual cells with time-lapse imaging.

**Fig. 4.**
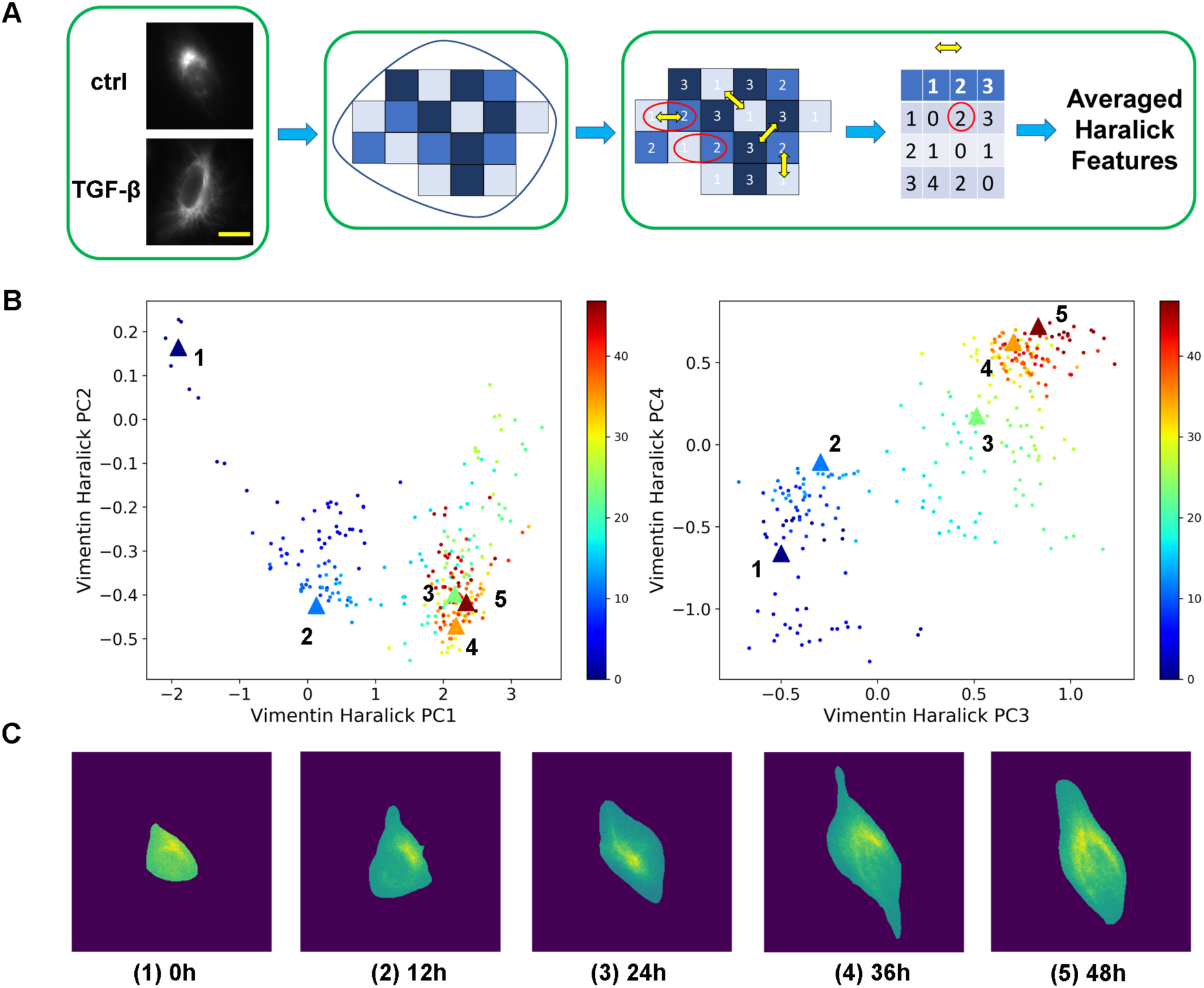
Haralick features quantify vimentin texture structures. (**A**) Flowchart of quantification. Left: typical vimentin fluorescence images of single cell before (top) and after treatment of 4 ng/ml TGF-β for two days (bottom, scale bar is 50 µm). Middle: Segmented single cell image with only pixels inside the cell mark kept for Haralick feature calculations. Right: Framework of calculating the single cell Haralick features. (**B**) Typical single cell trajectory on the plane of vimentin Haralick features PC1 and PC2 (left), and on the plane of PC3 and PC4 (right). Color bar represents time (units in hours). (**C**) Segmented single cell images of vimentin at various time points corresponding to the trajectory in panel B (labeled with large, triangular dots).

Texture features are widely used for image profiling in drug screening, phenotype discovery, and classification(*25–27*). We hypothesized that the texture features of vimentin can be quantified as an indicator of EMT progression. For quantification, we used Haralick features based on the co-occurrence distribution of grey levels. After segmentation, we calculated the grey level co-occurrence matrix (GLCM) in the mask of each cell, and 13 Haralick features based on the single cell GLCM and averaged all four directions (Fig. 4A, see Method for details). Nearly every Haralick feature shows a shift in distribution after TGF-β treatment (Fig. S4).

To capture the major variation in vimentin Haralick features during EMT, we performed linear dimension reduction with PCA (more details are in Method). Figures 4B and 4C show a typical single cell trajectory in the vimentin Haralick feature space, and corresponding segmented single cell images at various time-points along this trajectory, respectively. The dynamics in the vimentin space are again heterogeneous, as indicated here, and from two additional trajectories in Fig. S5.

### A label spreading function divides the combined morphological/texture space into three regions

Overall, we described a cell state in a 309 dimensional combined morphological/vimentin texture space. The cells occupy distinct regions in the morphological and vimentin texture features at the initial (0-2 hour) and final stages (46-48 hour) of 4 ng/ml TGF-β treatment (Fig. 5A). Physically, upon TGF-β treatment, the cell population relaxes from an initial stationary distribution in the morphological/vimentin space into a new one, and this study focuses on the dynamics of this relaxation process.

**Fig. 5.**
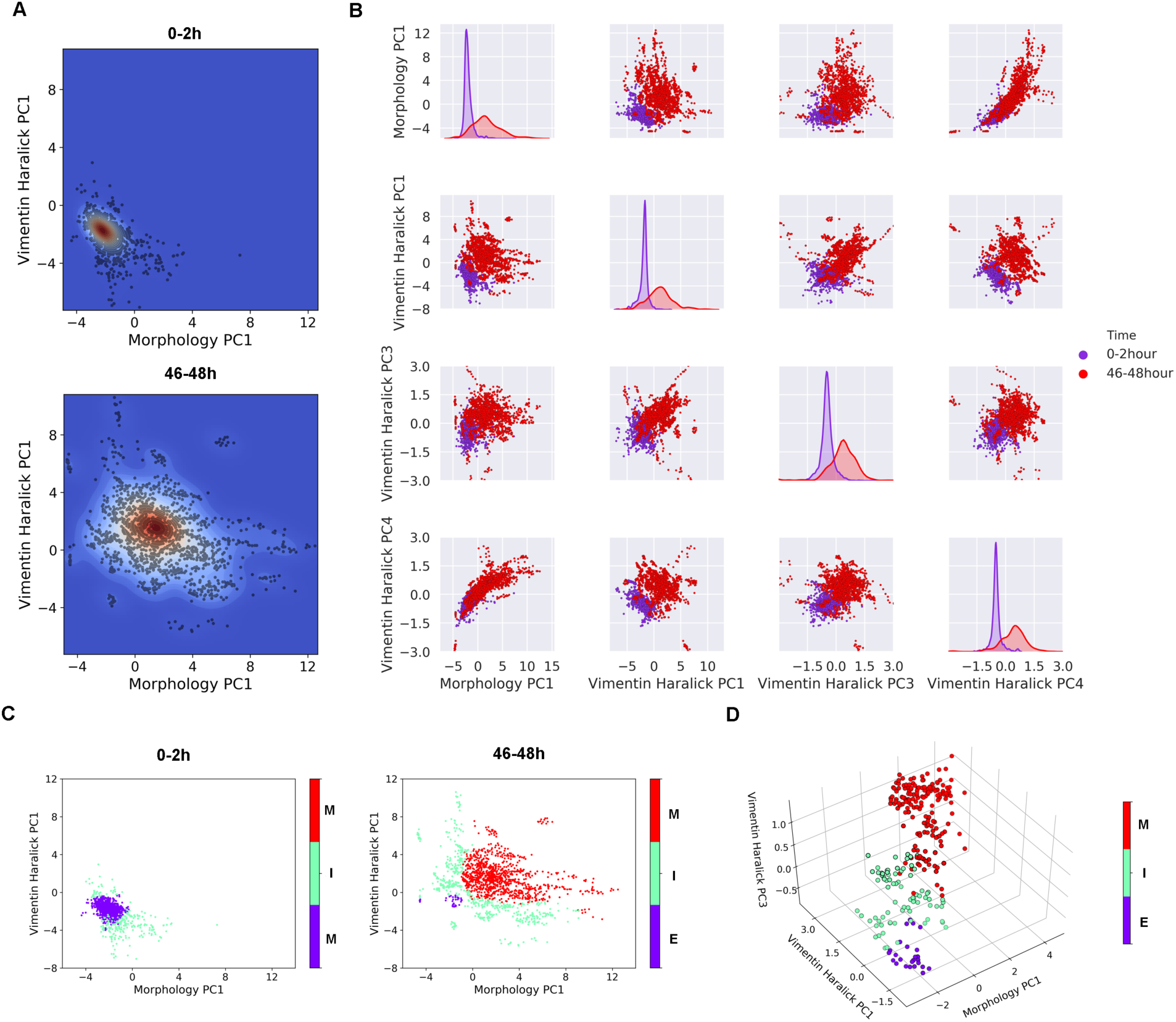
Single cell EMT states are defined in the morphology/vimentin feature space. (**A**) Kernel density plots in the plane of morphology PC1 and vimentin Haralick features PC1 estimated from measured single cell states (represented by dots), at 0 – 2 h (top) and 46 – 48 h after addition of TGF-β (bottom). (**B**) Distributions of cells at 0 – 2 h and 46 - 48 h cells after adding TGF-β in various features. The diagonal axes are plots of kernel density estimation of the 1D distribution of the corresponding features. (**C**) Scatter plot of 0-2 h data (left) and 46-48 h data (right) on plane of morphology PC1 and vimentin Haralick features PC1. Color represents cell state predicted by the fitted label spreading function. E: epithelial state, I: intermediate state, M: mesenchymal state. (**D**) A single cell trajectory with its state predicted by the label spreading function on the domain of morphology PC1, and vimentin Haralick features PC1 and PC3 (with states represented by different colors).

Close examination reveals major distribution shifts along four coordinates: morphology PC1 (82.7% variance of morphology), vimentin Haralick PC1 (57.0% variance), PC3 (8.9% variance) and PC4 (5.2% variance) (Fig. 5B, Fig. S6). This observation permits subsequent analyses restricted to these collective coordinates.

In the transition path theory or transition path sampling (*4, 16*), one divides the configuration space describing a reaction system into reactant, intermediate, and product regions. Specifically, for the present system, we developed a computational procedure that combines Gaussian mixture model (GMM) analysis of the cell distributions (Fig. S7A & B) followed by fitting a label spreading function (*28*) using the k-nearest-neighbor (KNN) method. The procedure divides the four-dimensional space into epithelial (*E,* or more precisely partial E for A549 VIM-RFP), intermediate (*I*), and mesenchymal (*M*) regions (Fig. 5C). Figure 5D shows a trajectory that starts within the *E* region, then progresses to the *I*, then the *M* regions.

### Single cell EMT trajectories follow distinct paths

According to the transition path theory, all the single cell trajectories similar to the one in Fig 5D (and Fig. S7C) that leave the *E* region, and end in the *M* region before returning to *E*, formed an ensemble of reactive trajectories. Overall, we recorded *N_T_* (= 196) acceptable continuous trajectories (see Methods), among them *N_R_* (= 139) are reactive trajectories (Movie S1 and Movie S2).

Single cell trajectories in the feature space show clear heterogeneous transition dynamics. In one representative trajectory (Fig. 6A, Fig. S8A left), the cell transits from the *E* to the *M* region following a series of transitions first along the vimentin Haralick PC1, then the morphology PC1. In contrary, in another trajectory (Fig. 6B, Figure S8A right) the cell proceeds with concerted morphological and vimentin Haralick feature changes.

**Fig. 6.**
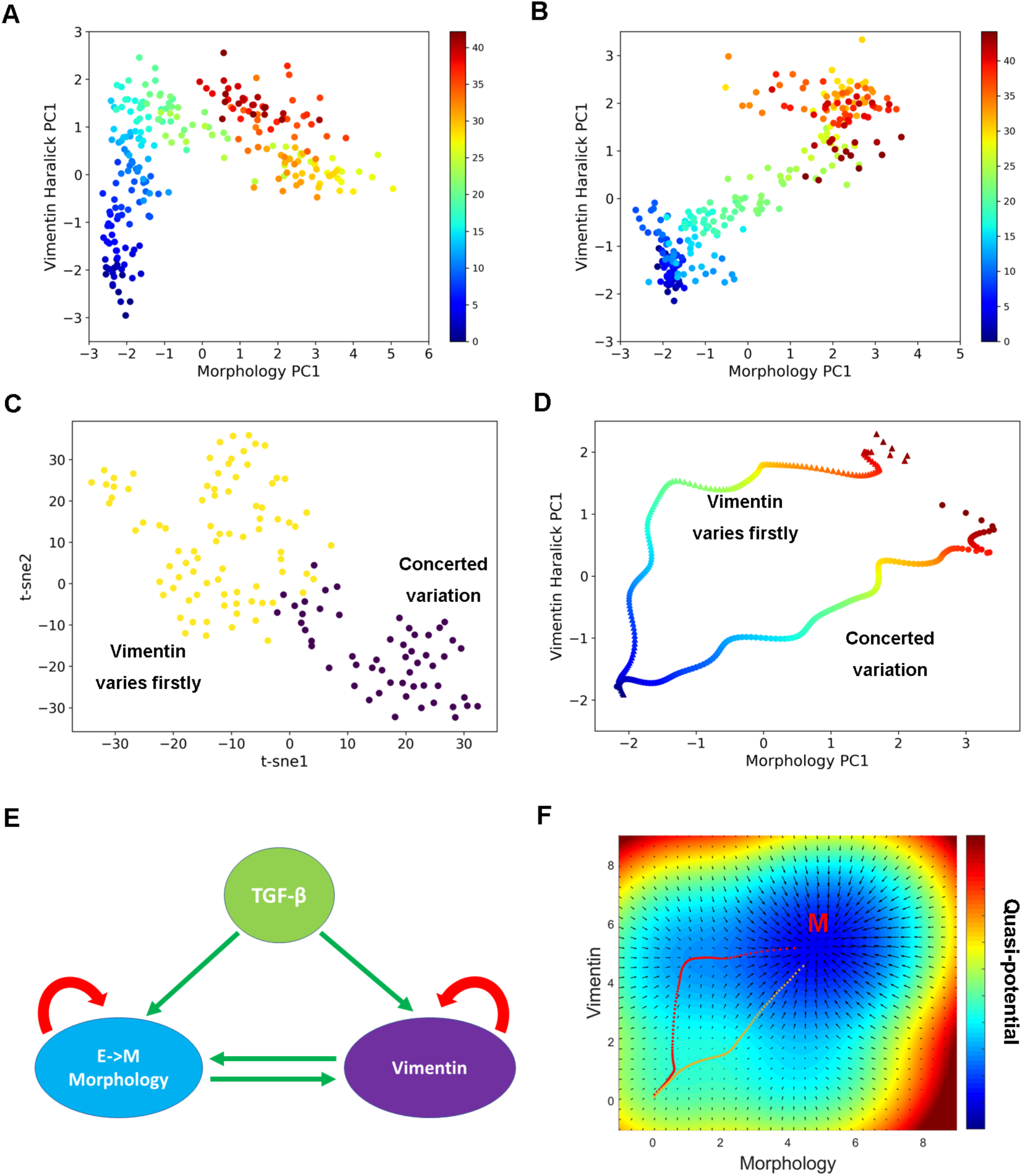
Single cell trajectory analyses reveal parallel paths of EMT. (**A**) A typical single cell trajectory in which the major change along the vimentin Haralick PC1 precedes the major change along the morphology PC1 (Class I). (**B**) A typical single cell trajectory in which the morphology PC1 and vimentin Haralick PC1 show concerted variation (Class II). (**C**) Projection of reactive trajectories on 2D t-SNE space using the DTW distances. Color represents labels of k-means clustering on the DTW distances. (**D**) Mean trajectories of Class I and II trajectories, respectively. They were calculated using the soft-DTW barycenter method. (**E**) Plausible mechanistic model. In this network, both morphology and vimentin changes are induced by TGF-β and they both activate each other and themselves. (**F**) Transition paths simulated with the model in panel E. The transition paths show dynamical characteristics similar to those observed experimentally.

To systematically study the two types of distinct behaviors, we used soft-dynamic time warping (DTW)(*29*) to calculate the distance between different trajectories, and t-SNE to project the trajectory distance matrix to 2D space(*30*). We found that these trajectories form two communities (Fig. 6C). A k-means clustering on these trajectories separated them into two groups, consistent with the two communities in the t-SNE space. The two groups of trajectories reveal different dynamical characteristics. In one group (Class I), vimentin Haralick PC1 varies firstly, followed by dramatic change of morphology PC1. In the other group (Class II), for most trajectories the morphology PC1 and vimentin Haralick PC1 change concertedly, while for only a small percentage the morphology PC1 change earlier than that of vimentin Haralick PC1 (Fig. S8B). Distinction between the two groups of trajectories is apparent from the scattered plot in the morphology PC1/vimentin Haralick PC1 plane (Fig. S8C), and the non-overlapping mean trajectories obtained using soft-dynamic time warping (DTW) barycenter(*29, 31*) (Fig. 6D and S8D).

To rule out the possibility that the existence of two classes of trajectories is an artifact of DTW, we analyzed cross correlation between morphology PC1 and vimentin Haralick PC1 of individual reactive trajectories. Cross-correlation analysis calculates the time delay at which the correlation between morphology PC1 and vimentin PC1 reaches a maximum value (*32*). The time delay shows a stretched distribution (Fig. S8E). A large portion of trajectories have vimentin Haralick PC1 change prior to morphology PC1 change, while another main group of trajectories have the time delay between morphology PC1 and vimentin Haralick PC1 close to zero. After separating the trajectories into two groups based on the sign of time delay, the mean trajectories of the two groups (Fig. S8E) are similar to what was obtained with k-means clustering on the DTW distance.

Therefore, a main conclusion of this study is that the live cell platform revealed two types of paths for the TGF-β induced EMT in A549 VIM-RFP cells. Figure 6E shows a plausible mechanistic model summarizing the existing literature. TGF-β activates morphological change and vimentin to induce EMT, while morphological change and vimentin expression can induce each other and themselves. More details of the model are in Materials and Methods. Computer simulations with the model followed by k-means clustering on DTW distance reproduce the two parallel EMT paths (Fig. 6F, Fig. S9C and Fig. S9D). The cross correlation analysis also showed results similar to what was observed experimentally (Fig. S9E). That is, the live-cell imaging platform presented here can provide mechanistic insight for further analyses.

## Discussion

Compared to the recent advances of fixed cell based single cell techniques, live cell imaging remains under-developed especially in studying CPTs due to some technical challenges. Generally speaking, the degrees of freedom specifying cell coordinates should be experimentally feasible for live cell measurement, and faithfully represent cell states. However, individual gene products typically only reflect partial dynamical information of a CPT process, and simultaneous fluorescence labeling of multiple genes is challenging. Recently, tracking cell morphological features through live cell imaging, emerges as a means of extracting temporal information about cellular processes in conjunction with expression-based cell state characterization(*11, 33–36*). Cellular and subcellular morphology reflects collective gene expression pattern and cell phenotype (*37, 38*). Furthermore, hundreds of or more morphology features such as cell size and shape can be conveniently extracted from bright field images without necessity of additional fluorescence labeling. Here, we further developed a quantitative framework for recording and analyzing single cell trajectories in a combined morphological/expression space, and a computational package for related image analyses.

Our application to the TGF-β induced A549 VIM-RFP EMT process demonstrates the importance of extracting dynamical information from live cell data. A cell has a large number of molecular species that form an intricately connected network, and it interacts with a fluctuating extracellular environment, including cell-cell interactions. Consequently, even iso-genetic cells show cell-to-cell heterogeneity, which further manifest as large trajectory-to-trajectory heterogeneity in single cell CPT dynamics, and some dynamical features characteristic to a particular process might be unavoidably concealed from snapshot data. Notably, our live cell data revealed information on the two distinct types of paths of EMT with distinct vimentin dynamics, in agreement with predictions from a mechanistic model based on previous reports that vimentin is a regulator as well as a marker of EMT.

For further mechanistic understanding beyond the phenomenological observation of the parallel paths, one needs to see how the cell expression pattern changes along the paths. One can use the paths identified from live-cell imaging data, rather than currently used pseudo-trajectory approaches based on a perceived expression-similarity criterion, to time-order snapshot single cell data, thus resolving the dilemma experienced in single cell studies. For this purpose, one needs to establish a mapping system between the morphological/texture space and the expression space. A better resolution of cell status in the morphological/texture space, such as the cell cycle stage of a cell, can reduce the observed cell-cell heterogeneity, and increase the fidelity of the mapping. Furthermore, within the present framework a better resolution of cell states in the morphological/texture space can be achieved by including additional features such as organelle texture and distributions in three-dimensions.

In summary, in this work we demonstrate that live cell imaging is necessary to reveal certain dynamical features of a CPT process concealed in snapshot data due to cell-cell heterogeneity. Meanwhile we present a framework that facilitates recent emerging efforts of using live-cell imaging to investigate how a CPT process proceeds along continuous paths at multiplex, albeit lower dimensional space, complementing to fixed-cell based approaches that can provide snapshots of genome-wide expression profiles of individual cells. We expect that the framework be generally applied since dramatic morphological changes typically accompany a CPT process.

## Materials and Methods

### Cell Culture and Treatment

The human non-small cell lung carcinoma line, A549 (ATCC CCL-185) and A549 VIM-RFP (ATCC CCL-185EMT) were from American Type Culture Collection (ATCC). Cells were cultured in F-12K medium (Corning) with 10% Fetal Bovine Serum (FBS) in Mattec glass bottom culture dishes (P35G-0-10-C) in a humidified atmosphere at 37°C and 5% CO_2_. Culture medium was changed every 3-5 days. During imaging, Antibiotic-Antimycotic (100X) (Thermo Fisher 15240062) and 10 mM HEPES (Thermo Fisher 15630080) was added to the culture medium.

### sgRNA design and cloning into the gRNA-expression vector

The CHOPCHOP website (https://chopchop.rc.fas.harvard.edu/) was used to design high-performance sgRNAs to target the sequence near the stop codon of the human vimentin gene. The cleavage activities of the gRNAs were validated using the T7E1 assay according to the manufacturer’s instructions (NEB, #E3321). The sgRNA VIM-AS3 (5’-CTAAATTATCCTATATATCA-3’) was chosen in this study. To generate the sgRNA expressing vector, VIM-AS-3 gRNA oligos were designed, phosphorylated, annealed, and cloned into the PX458 (Addgene catalog no. 48138) vector, using BbsI ligation. Multiple colonies were chosen for Sanger sequencing to identify the correct clones using the primer U6 Fwd: 5’-AAGTAATAATTTCTTGGGTAGTTTGCAG-3’

### Construction of VIM-RFP knock-in donor

The VIM-RFP knock-in donor was designed and constructed to contain approximately 800 bp left and right homology arms, a Cayenne RFP gene (Atum #FPB-55-609), preceded by a 22 amino acids linker, and followed by a bovine growth hormone polyadenylation signal sequence. To assist in drug-based selection of gene edited cell clones, an EF1α-blasticidin selection cassette, flanked by Loxp sites, was also cloned into the vector, and positioned upstream of the right homology arm.

### Generation of A549 VIM-RFP reporter cell clones

CRIPSR/Cas9 technology was utilized to incorporate the RFP reporter into the 3’ terminal end of the vimentin gene. Briefly, A549 cells were plated at a density of 2 x 10^5^ cells/well in a 6-well plate. After 24 hours, cells were transfected with 4.0 ug PX458_VIM-AS3 plasmid, 4.0 ug VIM-RFP knock-in donor plasmid, and 24 ul transfeX (ATCC ACS-4005). Blasticidin selection (10 ug/ml) was applied 24 hours post-transfection. RFP positive cells were single cell sorted and expanded for molecular characterization.

### VIM-RFP knock-in clone identification and confirmation

RFP positive A549 VIM-RFP cells were harvested and DNA was extracted using QuickExtract (Epicentre, QE09050). Primers were designed for left homology arm and right homology arm junction PCR (left junction Fwd: 5’-TAGAAACTAATCTGGATTCACTCCCTCTG-3’, left junction Rev: 5’-ATGAAGGAGGTAGCCAGGATGTCG-3’; right homology Fwd: 5’-ATTGCTGCCCTCTGGTTATGTGTG-3’, right homology Rev: 5’-ATTACACCTACAGTTAGCACCATGCG-3’); Junction PCR was performed using Phire Hot Start II DNA Polymerase (Thermo Scientific), and the PCR amplicons were subjected to Sanger sequencing for identification of clones that contained the expected junction sequences at both left and right homology junctions.

### Immunostaining

A549 VIM-RFP cells were washed with PBS, fixed with 4% formaldehyde, and blocked with 5% normal goat serum /0.1% Triton X-100 in PBS for 30 mins. Afterwards, the primary antibodies were added to the blocking buffer and cells were incubated for 1 hour at room temperature. Cells were subsequently washed and incubated with the secondary antibodies for 1 hour, wrapped in aluminum foil. After washing, cells were covered by 50% glycerol and images were taken with a Nikon Ti-E microscope (Hamamatsu Flash 4.0 V2). The primary antibodies were mouse anti-N-Cadherin (13A9) (Cell Signaling Technologies, Cat#14215) (1:100 dilution) and mouse anti-Snail (L70G2) (Cell Signaling Technologies, Cat#3895) (1:300 dilution). For secondary antibodies, goat anti-mouse Alexa Fluor 647 (Thermo Scientific, Cat #A-21235) was used at a 1:1000 dilution.

### EMT Induction

A549 VIM-RFP cells were plated at a density of 1x10^4^ cells/cm^2^ and maintained in F-12K medium (ATCC 30-2004) supplemented with 10% FBS (ATCC 30-2020). After 24-48 hours, culture medium was replaced with fresh medium supplemented with 4.0 ng/ml TGF-β (R&D Systems 240-B) for 1-3 days to induce EMT. Non-treated cells were used as a control.

### Matrigel invasion assay

Control and EMT induced A549 VIM-RFP cells were seeded into inserts of Boyden chambers (BD Biosciences, San Jose, CA) that were pre-coated with Matrigel (1mg/ml), at 5x10^4^ cells per insert in culture medium without FBS, and then inserts were transferred to wells with culture medium containing 10% fetal bovine serum as a nutritional attractor. After 24 hours incubation, invading cells on the bottom side of the insert membrane were fixed with 4% paraformaldehyde for 2 min, permeabilized with 100% methanol for 20 min, and stained with 0.05% crystal violet for 15 min at 37°C. Non-invading cells on the top side of the membrane were removed by cotton swab. Photographs were taken from five random fields per insert. Cells in the five random fields were counted.

### Western blot analyses

A549 parental and VIM-RFP cells were harvested and lysed in ice-cold RIPA buffer containing protease inhibitors, followed by sonication and centrifugation. Supernatant was taken for protein quantification using the Pierce BCA protein assay kit (Thermo Fisher, cat# 23227). 10µg of protein was loaded onto a 4-20% Bio-Rad SDS mini-protean TGX gel, which was run at 130V for approximately 1 hour. The protein was then transferred to a PVDF membrane using the Bio-Rad wet transfer system at 30V for 2 hours. The membrane was blocked for 1 hour with 5% milk in TBS-T, then incubated with anti-vimentin antibody (clone D21H3, Cell Signaling Technologies, (1:500 dilution in 5% milk/TBS-T) and with anti-GAPDH antibody (Abcam, AB37168, 1µg/mL in 5% milk/TBS-T) overnight at 4°C. The blots were then washed 3 x 15 minutes in TBS-T followed by secondary antibody incubation using goat anti-rabbit poly clonal horse radish peroxidase (HRP) conjugated secondary antibody (1:10,000 dilution in 5% milk/TBS-T) for 1 hour at room temperature. The blots were then washed again 3 x15 minutes and treated with Bio-Rad Clarity ECL for 5 minutes. The blots were developed using a BioRad Gel Doc XR+ imaging system (BioRad 1708195).

### Off Target analysis

The top 10 potential off-target sites against VIM-AS3 gRNA identified by the CHOPCHOP website (https://chopchop.rc.fas.harvard.edu/) were used to evaluate the off-target cleavage in A549 VIM-RFP cells. PCR primers were designed to span each of the 10 mis-match off-target sequences. PCR amplicons were sequenced and mis-match sequences were analyzed for DNA cleavage.

### Imaging

Time-lapse images were taken with a Nikon Ti-E microscope (Hamamatsu Flash 4.0 V2) with differential interference contrast (DIC) and Tritc channels (Excitation wavelength is 555 nm and Emission wavelength is 587) (20 × objective, N.A. = 0.75). The cell culture condition was maintained with Tokai Hit Microscope Stage Top Incubator. Cells were imaged every 5 min with the DIC channel and every 10 min with the Tritc channel. The exposure time for DIC was 100 ms and the exposure time for the Tritc channel was 30 ms. That is, each full (two-day long) single cell trajectory contains 577 DIC images and 289 fluorescent images.

### Single cell segmentation and tracking

We segmented single cells using a previously developed method combining deep convolution neural networks (DCNN) and watershed(*39*). To quantify cell morphology, we adopted the active shape model method(*17, 20*). After single cell segmentation, the cell outline was extracted and resampled into 150 points. All the single cell outlines were aligned to a reference outline (calculated based on the average of several hundred cells). The 150 points (x and y coordinates) are the 300 features of cell morphology.

For single cell tracking, we used the *TrackObjects* module in CellProfiler on the segmented images using a linear assignment algorithm (*40, 41*). In long-term imaging, the accurate tracking of cells can be lost for several reasons, such as cells moving in or out of the field of view, or inaccurate segmentation. We kept trajectories that were continuously tracked with the starting point no later than 12 hours and the end point no earlier than 30 hours after adding TGF-β. These 196 trajectories were used for subsequent principal component analysis. Among them, 139 were identified as reactive trajectories.

### Vimentin image analysis

Haralick features have been widely used for classifying normal and tumor cells in the lungs (*42*), and the subcellular features or patterns such as protein subcellular locations (*43, 44*). After cell segmentation, each cell was extracted and its Haralick features were calculated using mahotas (*45*). Haralick features describe the texture as coarse or smooth, and complexity of images(13 features)(*43*). Haralick feature calculation was based on the grey level co-occurrence matrices (GLCM) (*46*). The GLCM’s size was determined by the number of grey levels in the cell image.

Due to cell heterogeneity, the numbers of grey levels varied in different cells. Because the GLCM has four directions (0,45, 90 and 135 degrees), the Haralick features were averaged on all four directions to keep rotation invariance.

### Scaling of single cell trajectories

Due to cell heterogeneity, it is more informative to examine the temporal change of an individual cell relative to its initial state, such as the basal level of gene expression in signal transduction studies(*10, 47, 48*). For the present system, it is the initial position in the combined morphology/texture space. We used a stay point searching algorithm(*49*) to find the initial stay point of each cell in the space of morphology and vimentin Haralick features. For each trajectory, we scaled all the landmark points by the square root of the area of the initial stay point. Physically, the latter is a characteristic length of the cell, and the scaling reflects the observation that the cell size does not affect EMT(*50*). All the vimentin Haralick features were reset so that the values at the initial stay point assume zero. The principal components were calculated after scaling. The scaling allows one to examine the relative temporal variation of single cells.

Principal component analysis (PCA) was performed on all *N_T_* trajectories, i.e., a total of *N_c_* (49689 cells) with 300 morphology features (*N_c_* × 300 matrix) for linear dimensionality reduction(*51*). The first seven components explained more than 98% of the variance. Specifically, the first and second components explained 82.7% and 10.5% of the variance, respectively. After calculation of Haralick features for each cell, PCA was calculated on the *N_c_*× 13 matrix for linear dimension reduction.

### Procedure of defining regions in the morphology/texture feature space

We fitted the distribution on each of the four morphology/texture coordinates with a two-component (*c*_1_, *c*_2_) Gaussian mixture model (GMM) separately (Fig. S7A)(*51*), and used the four GMMs to define the *E*, *I*, and *M* states (Fig. S7B).

For each single cell in the space of morphology PC1, vimentin Haralick PC1, PC3 and PC4 *X* (*x_i_* | *i* = 1, 2,3, 4), the label of each coordinate *L_i_* is defined using the GMM with the following equations:

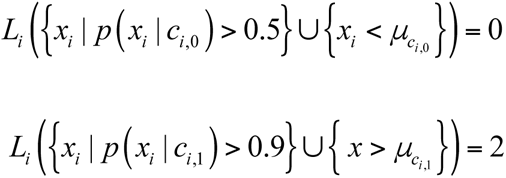

Where *p* (*x_i_* | *c_i_*, …) is the posterior probability of certain component of 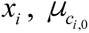 and 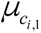 are the mean values of the two components of *i* − *th* GMM. The label of complementary set is defined as 1.

With the labels defined on all four coordinates, we first defined the *E* state as 
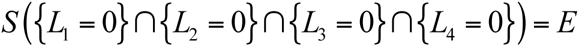

the *M* state as 
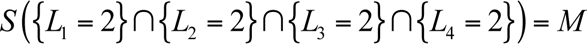

and the *I* state otherwise. However, this definition suffers from one weakness in that it assigns the same weight to vimentin Haralick PC1, PC3 and PC4, although vimentin Haralick PC3 and PC4 count for less variance than the PC1 does. Because of this equal weight distribution, small fluctuations in vimentin Haralick PC3 and PC4 could lead to unstable assignment of cell states.

To solve this problem, we use the above definition as an initial estimate of cell state to fit to a label spreading function(*28, 51*). When fitting the label spreading function, we adopted the k-nearest-neighbor (KNN) method (50 neighbors) and use a high clamping factor (0.5) to assure the global and local consistency. The KNN algorithm in the label spreading function allows one to take the different scales of vimentin Haralick PC1, PC3, and PC4, and community structure into consideration (Fig. 5C). Since PC1 is more important in defining the range of neighbors, the weight of PC1 in the definition was automatically increased. This change in definition avoids the situation that cells belonging to a common community and close to each other in the morphological/feature space, get assigned to different cell states.

### Cross correlation analysis

The cross correlation was used to calculate the time delay between different signals. We observed different transition time of morphology PC1 and vimentin Haralick PC1. The cross correlation of the two time series (of morphology PC1 and vimentin Haralick PC1) were calculated(*52*). The time lag between the two signals was set for when the value of cross correlation reaches the maximum(*32*). We separated all the trajectories by the sign of time delay between morphology PC1 and vimentin Haralick PC1.

### Stochastic simulation on double well potential (Fig. 1)

While a cellular system is far from thermodynamic equilibrium, for simplicity we illustrated the effect of cell-cell heterogeneity due to hidden slow variables with the following model system (and the computer code is included as a supplemental file), The potential function is *U* = ((*o* − 2*h*)^2^ − 2) + 3*h*^4^, where *o* is the observable and *h* is the hidden slow variable (Fig. 1A).

The simulations were performed using the following procedures:

1. Generate initial condition (*o*_0_, *h*_0_) in the left well of this potential of multiple trajectories (4581) (Fig. S1A) with the Metropolis-Hastings algorithm(*53*).
2. For each initial condition, with fixed hidden slow variable *h*_0_, propagate the observable *o* with Langevin simulations along the 1D potential and with a Gaussian white noise (*η*(*t*)). 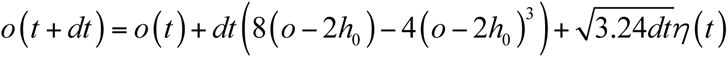, where *^dt^* is 0.01(Figure S2B shows an example 1D potential with *h*_0_ = 0.2).
3. Propagate each trajectory to t = 10.

### Network model of EMT (Fig. 6E)

We built the network model by summarizing the existing literature. The EMT morphology variation is mainly contributed by generation of filament actin and E-cadherin down-regulation, which can activate the YAP/TAZ pathway (*54*). YAP/TAZ pathway can induce translocation of Smad2, which play important roles in EMT(*55, 56*). Thus, morphology variation can activate both vimentin and itself. Vimentin can induce the EMT morphological change through regulating β1-intergrin and E-cadherin(*57*). Vimentin can be activated upon TGF-β induction through Slug and it also activate Slug through dephosphorylation of ERK, which forms a self-activation loop (*58*). Vimentin is required for the mediation of Slug and Axl (*57, 59–61*), and it can induce variation of cell morphology, motility and adhesion (*61*). Vimentin fibers regulate cytoskeleton architecture (*57*), and more vimentin fibers are assembled in A549 cells during EMT (*22*).

Next we formulated a mathematical model corresponding to the network:

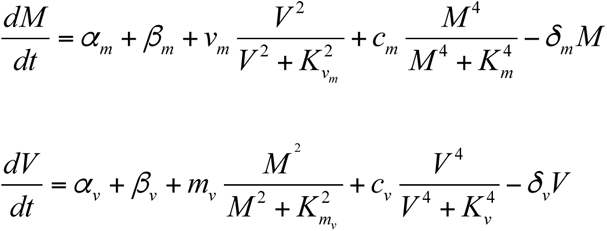

Where *α_m_* (= 0.1) and *α_v_* (= 0.12) are the basal generate rates of morphology variable and vimentin separately, *β_m_* and *β_v_* are the generate rates of morphology change and vimentin activated by TGF-β, respectively, *v_m_* (= 2.0) and *m_v_* (= 1.0) are the activation coefficients of vimentin and morphology to each other, *c_m_* (= 3.8) and *c_v_* (= 4.0) are the self-activation coefficients of morphology and vimentin, respectively, and *K*_v_m__ (= 8.0), *K*_m_v__ (= 8.0), *K*_v_ (= 2.4) and *K_m_* (= 2.4) are the half maximal effective concentrations of the Hill function. *δ*_m_ (1.0) and *δ_v_* = 1.0) are degradation rates of morphology and vimentin, respectively.

The Langevin simulations were performed as follows:

1. Set *β_m_* and *β_v_* as 0 for simulating the control condition (i.e., without TGF-β treatment). Initialize a trajectory with random point (*M*_0_, *V*_0_) sampled from a uniform distribution within a range of [0, 5) . Run simulations with the following equation: 

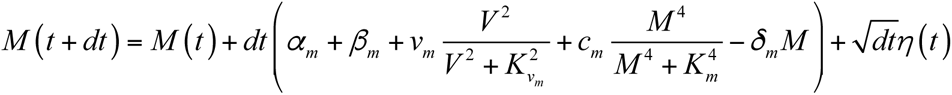

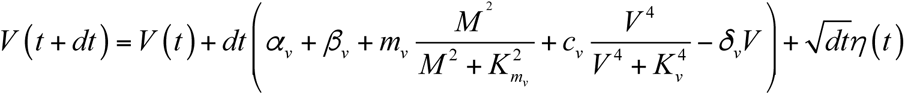 where *dt* was set to be 0.01, and *η* (*t*) was normal Gaussian white noise. The duration of simulation was set to 100. At the end of the simulation, the cell state relaxed to the basin of the epithelial state (Fig. S9A, which was then set as the initial condition under TGF-β treatment.
2. After generating multiple initial conditions from the first step, increase the values of *β_m_* (to 0.6) and *β_v_* (to 1.0) to simulate the condition of TGF-β treatment. If the cell gets into the range that its distance to the attractor of the mesenchymal state (Fig. 6F and Fig. S9B) is less than one, this trajectory was considered as a trajectory of EMT.
3. After getting *N_simu_* (= 185) reactive EMT trajectories, we performed analysis with the simulated trajectories similar to we did with the experimental trajectories.

We obtained the steady state probability distribution *P_ss_* by solving the diffusion equation (using *Matlab 2018a PDEtool*), 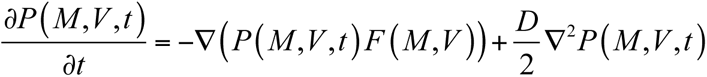 where 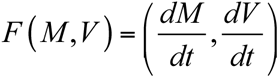 and *D* is the diffusion coefficient. With the steady state probability distribution, we obtained the quasi-potential of EMT defined as *U* ∝ − ln (*P_ss_*) (*62, 63*). Without TGF-β, there exists a deep basin as epithelial state and a shallow basin as mesenchymal state in the quasi-potential landscape (Fig. S9A). After TGF-β treatment, the landscape is changed, on which the mesenchymal basin becomes deep, and a valley is formed where the vimentin level is high (Fig. S9 B and Fig. 6E).

### Computer package M-TRACK

The M-track program is written in Python 3 and provided with a graphical user interface (GUI). It provides tools for analyses of cell morphology with the active shape model, distribution and texture features of protein or gene florescence in single cell, and single cell trajectories in the PC domain. The input files include the original grey-level images, segmented cell mask and database file of tracking results from Cellprofiler. The computer package can be downloaded from GitHub (https://github.com/opnumten/M-TRACK). Part of the source code is adapted from Celltool(*20*).

### Statistical Analysis

Statistical analyses were performed mainly with Python package including Scipy and Scikit-learn(*51, 52*). Student ‘s t-test was used to calculate the statistical difference between different groups of samples. The samples for imaging were randomly selected to avoid bias.

## Acknowledgments

**General**: We would like to thank Drs Yi Jiang and Xiuxiu He on helpful discussions of the active shape model.

## Funding

This work was partially supported by the National Science Foundation [DMS-1462049], National Cancer Institute [R01CA232209], National Institute of Diabetes and Digestive and Kidney Diseases (R01DK119232) to JX, and the NIH supported microscopy resources in the Center for Biologic Imaging at University of Pittsburgh (1S10OD019973-01).

## Author contributions

JX conceived the project, D.D., S.K., M.E.M., and W.S. generated and characterized the cell line, W.W. performed experiments with support from C.T.W. and S.C.W. on imaging, and from J.Z., Y.J.C., and Y.Y.C. on cell culturing, W.W. and J.X. analyzed the data with input from Y.J.C. and Y.Y.C., Y.D. generated the GUI of the computer package, W.W. and J.X. wrote the manuscript with input from J.Z., S.C.W., D.D., and W.S.

## Competing interests

D.D., S.K., M.E.M., and W.S. are employees of the American Type Culture Collection, which offers the A-549 VIM RFP cell line for sale commercially.

## Data and materials availability

The computer package together with the image data files can be downloaded from GitHub (https://github.com/opnumten/M-TRACK).

## Supplementary Materials

**Figure S1.**
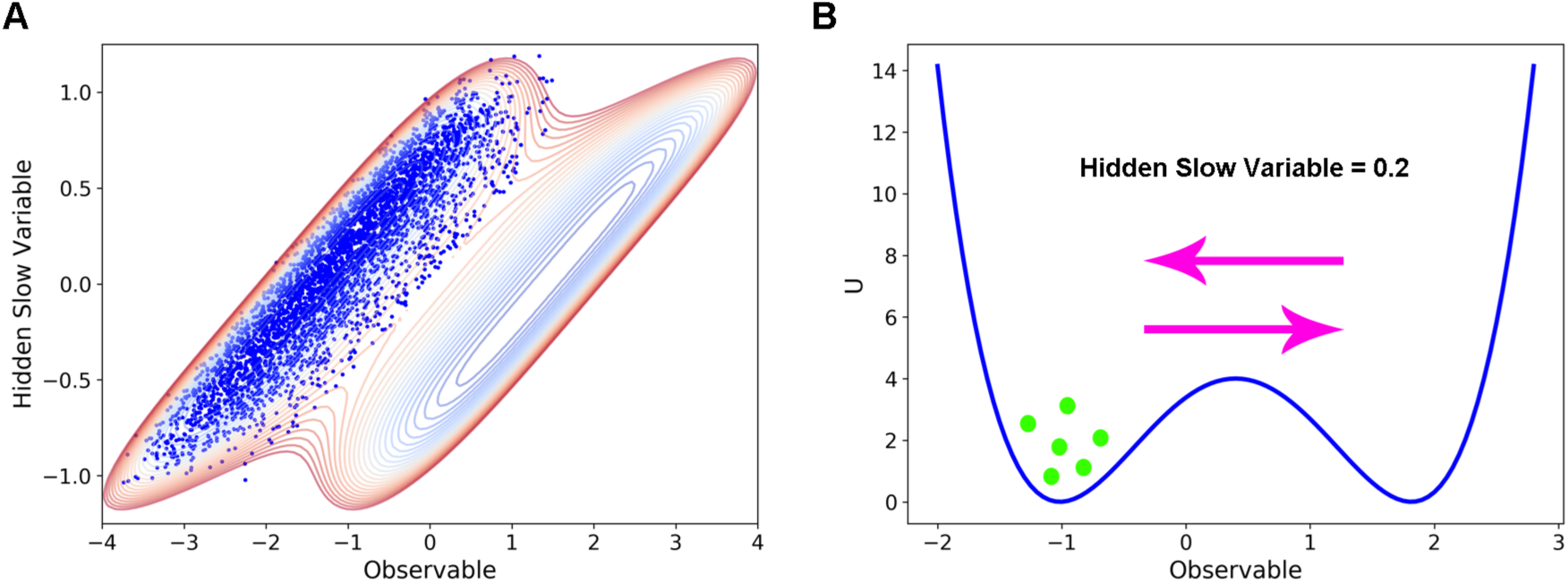
Additional details for the potential system simulations. (**A**) Metropolis-Hastings sampling of the initial conditions of the left well (blue dots). Also shown is the contour plot of the potential system. (**B**) A 1D potential slice with the hidden slow variable set to be 0.2. All the simulations of single trajectories start from the left well and jump between two wells for a fixed duration. Single trajectory can end in either well.

**Figure S2.**
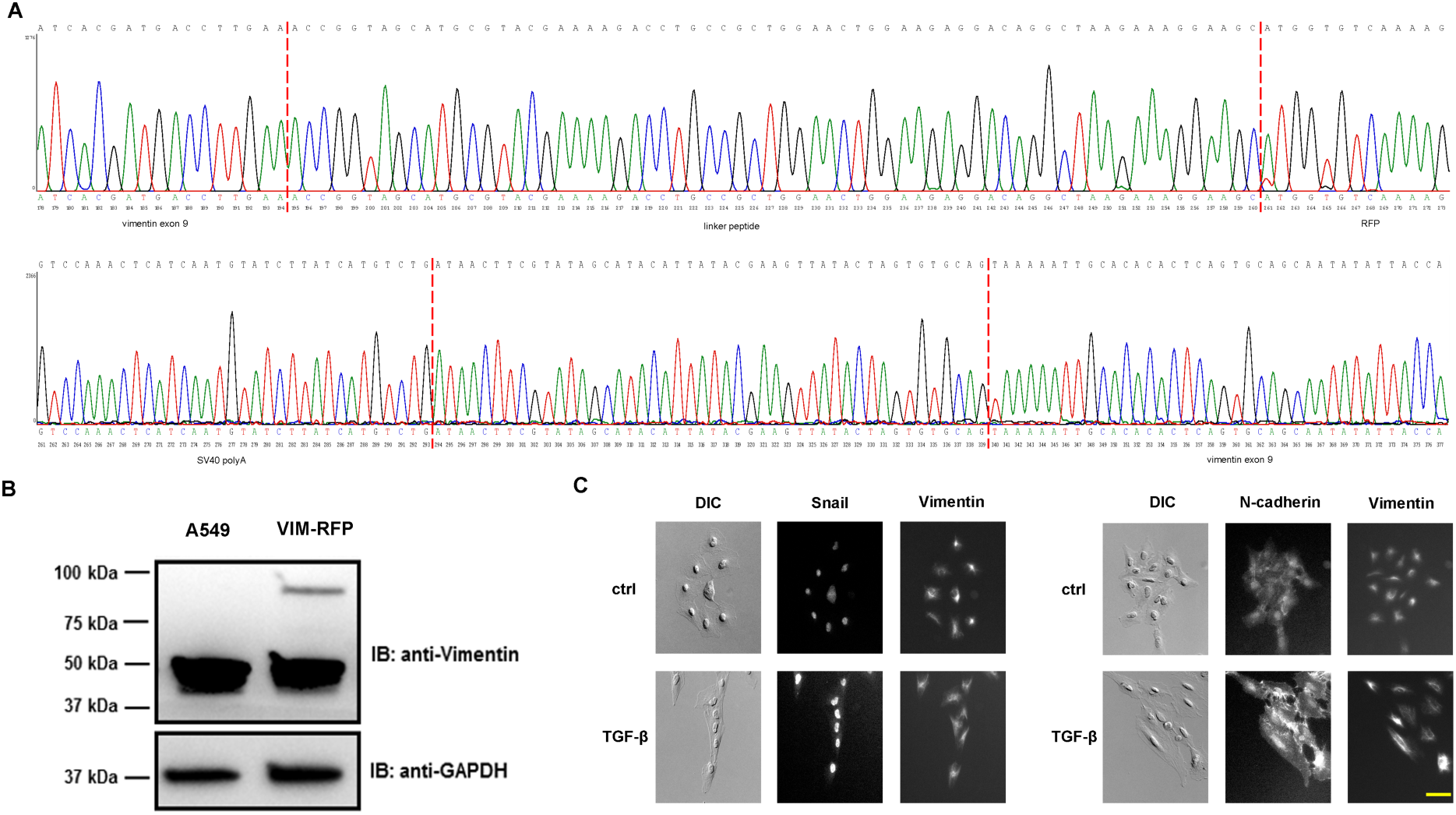
Confirmation and characterization of the knock-in for the A549 vimentin-RFP cell line. (**A**) Sequencing result of the knock-in for the A549 vimentin-RFP cell line. (**B**) Western blot of the knock-in for the A549 vimentin-RFP cell line. (**C**) Immuno-staining images of Snail and N-cadherin together with cell shape and vimentin variation after TGF-β induction. After TGF-β induction, the expression level of snail and N-cadherin increased along with variation of morphology and vimentin. Scale bar is 50 µm.

**Figure S3.**
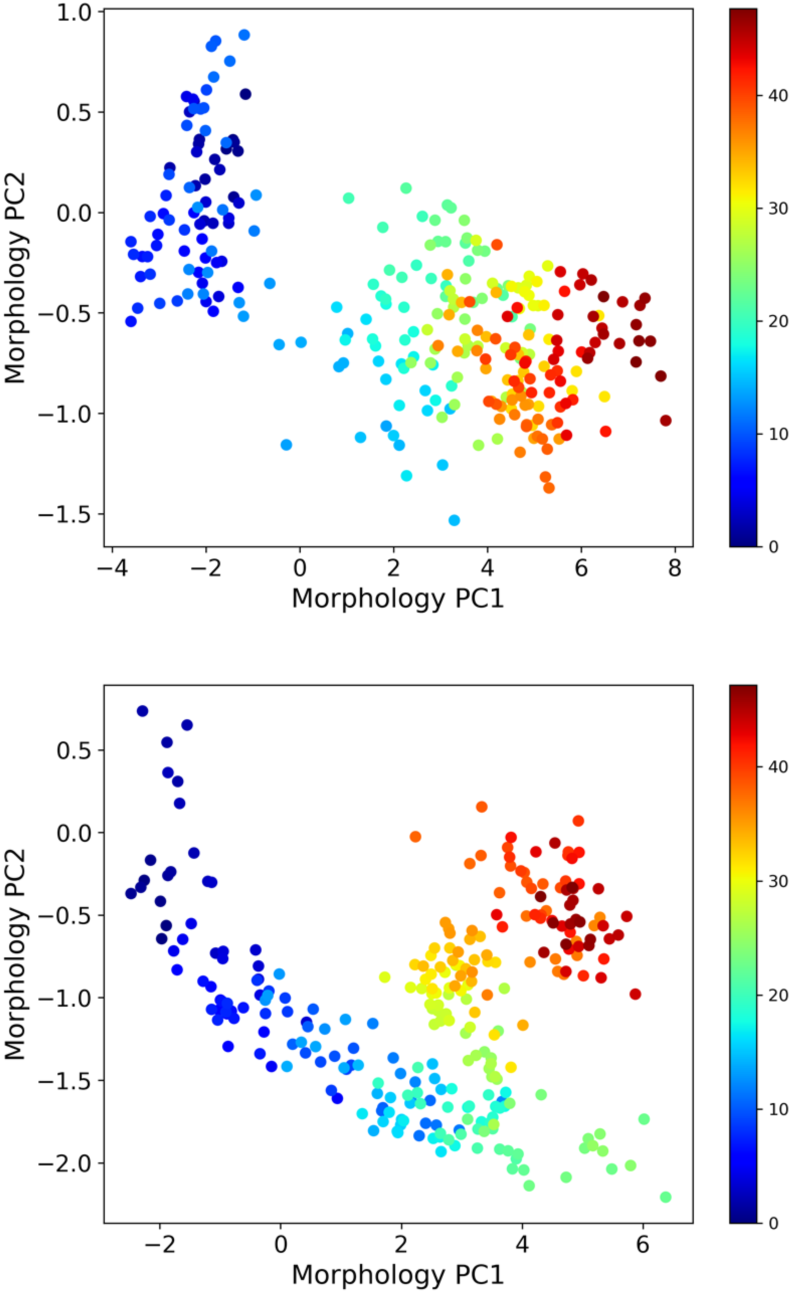
Additional examples of single cell trajectories in the morphology PC domain.

**Figure S4.**
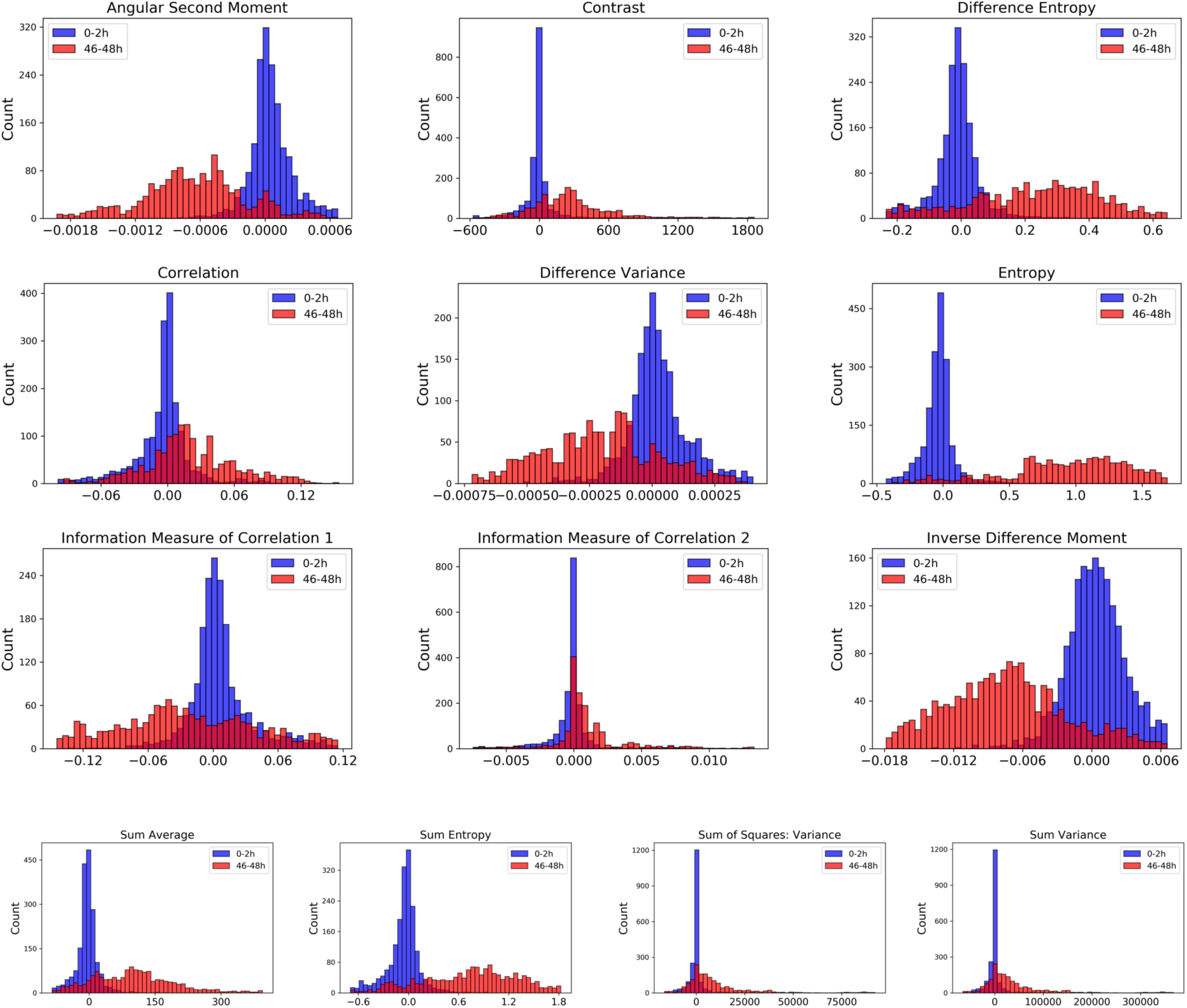
Distributions of Haralick features of cells with and without TGF-β treatment. Blue and red color represent cells 0-2 hours and 46-48 hours after adding TGF-β, respectively.

**Figure S5.**
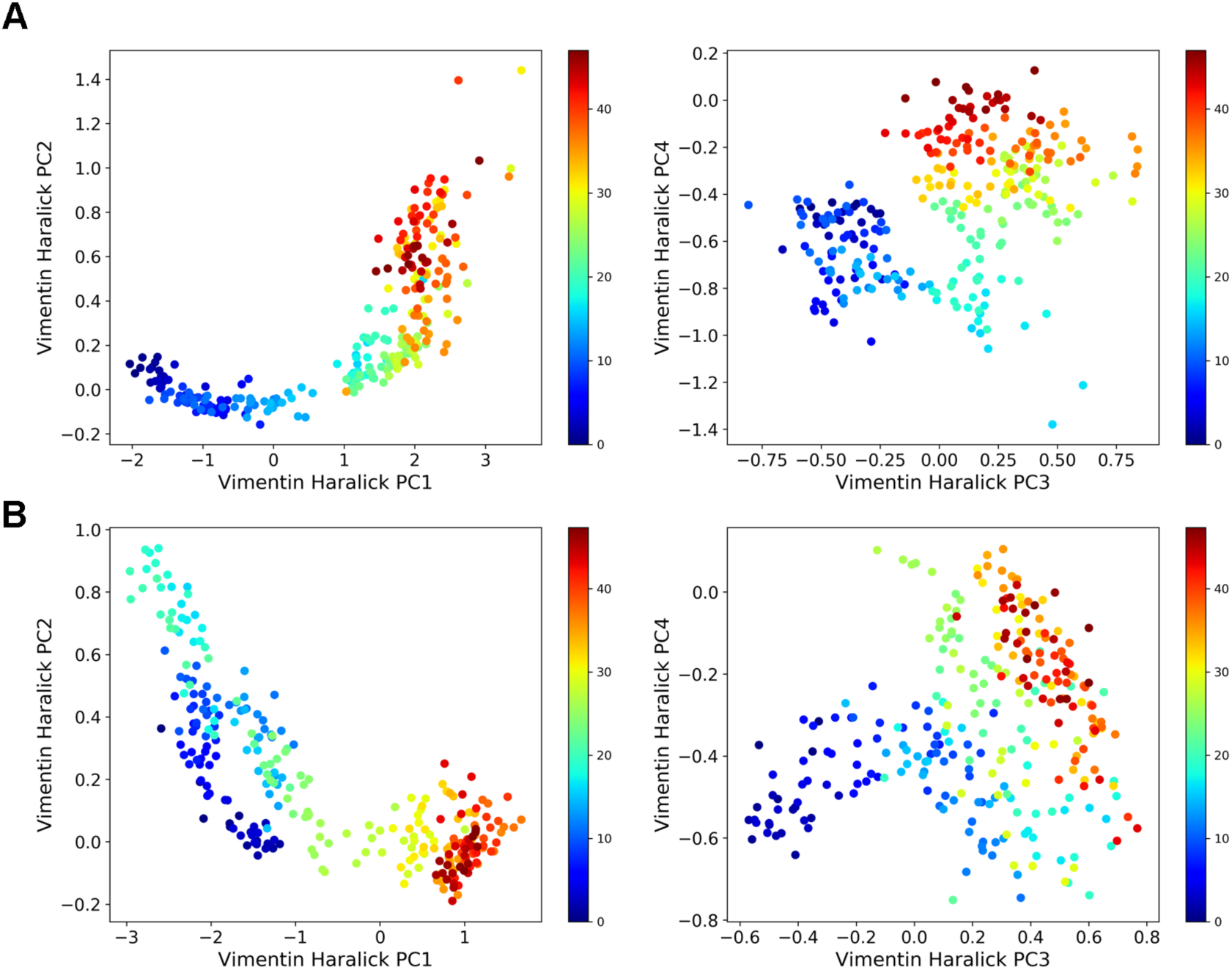
Additional examples of single cell trajectories. The trajectories are plotted in the plane of vimentin Haralick PC1-PC2 (left) and vimentin Haralick PC3-PC4 (right), respectively.

**Figure S6.**
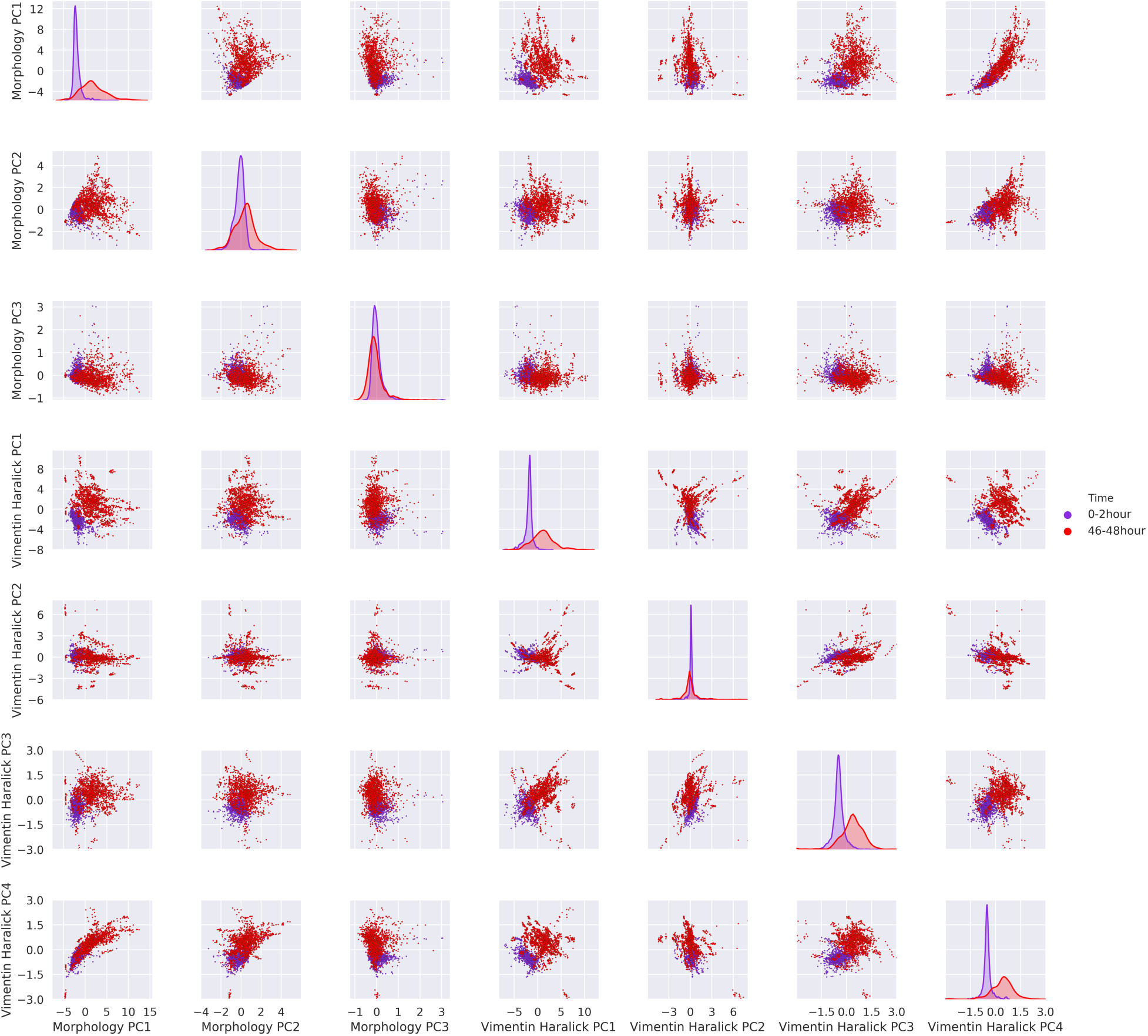
Distributions of various cellular features of cells. Purple and blue represent results at 0 – 2 hours and 46 - 48 hours after adding TGF-β, respectively. The diagonal axes are plots of kernel density estimation of the 1D distribution of corresponding features.

**Figure S7.**
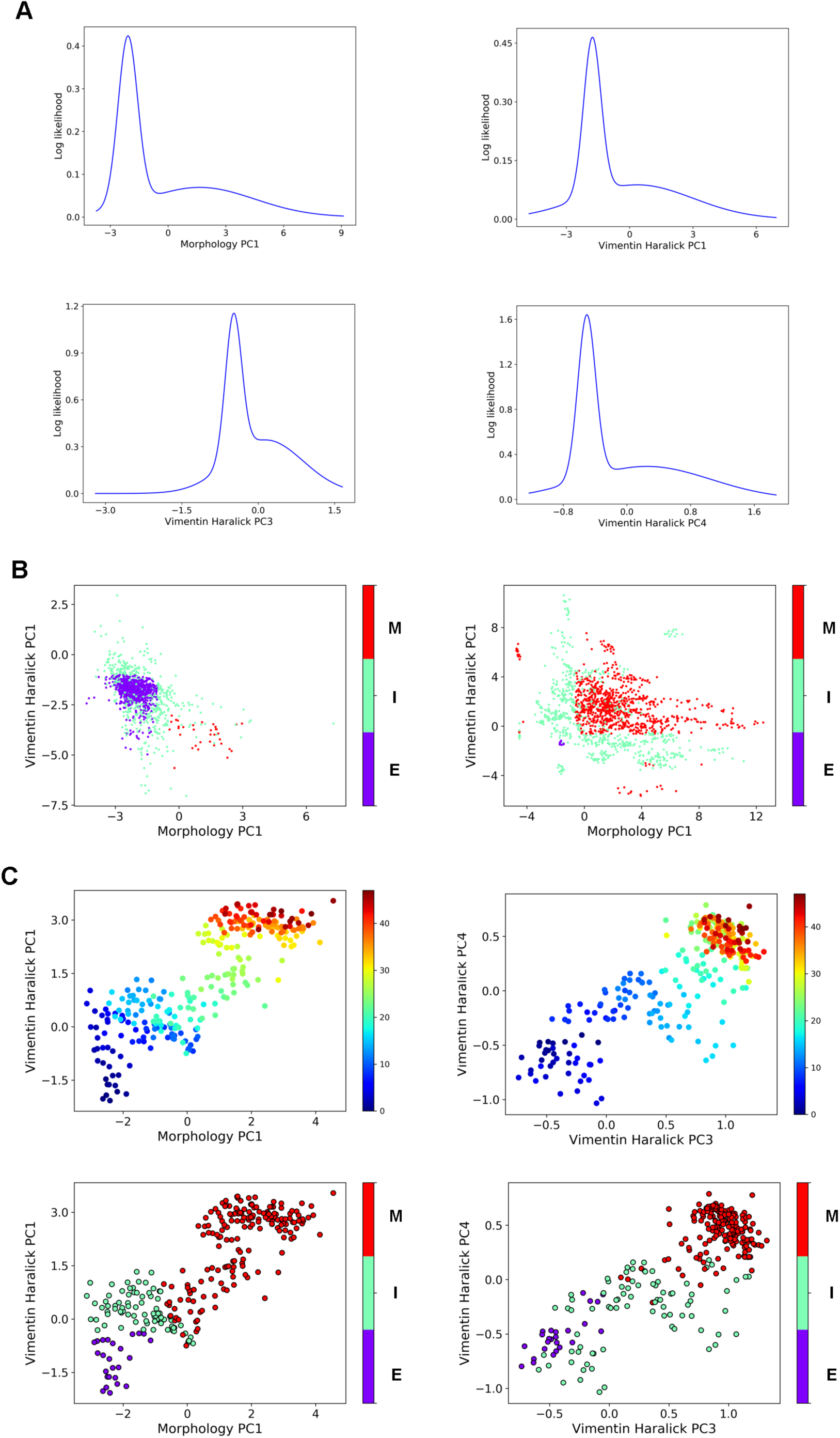
Additional results for defining cell states. (**A**) Log likelihood plots of fitted GMMs for cell feature distributions of cells 0-2 hours and 46-46 hours after adding TGF-β. (**B**) Scatter plots of initial estimate of states of cells at 0-2 hours (top) and 46-48 hours (bottom) on the plane of morphology PC1 and vimentin Haralick features PC1 defined by GMMs, which were fitted on morphology PC1, vimentin Haralick features PC1, PC3, and PC4 separately (color represents cell state). (**C**) The single cell trajectory in Fig. 5D (top, color represents time in hours) and its state, predicted by the label spreading function in various representations (bottom, color represents state).

**Figure S8.**
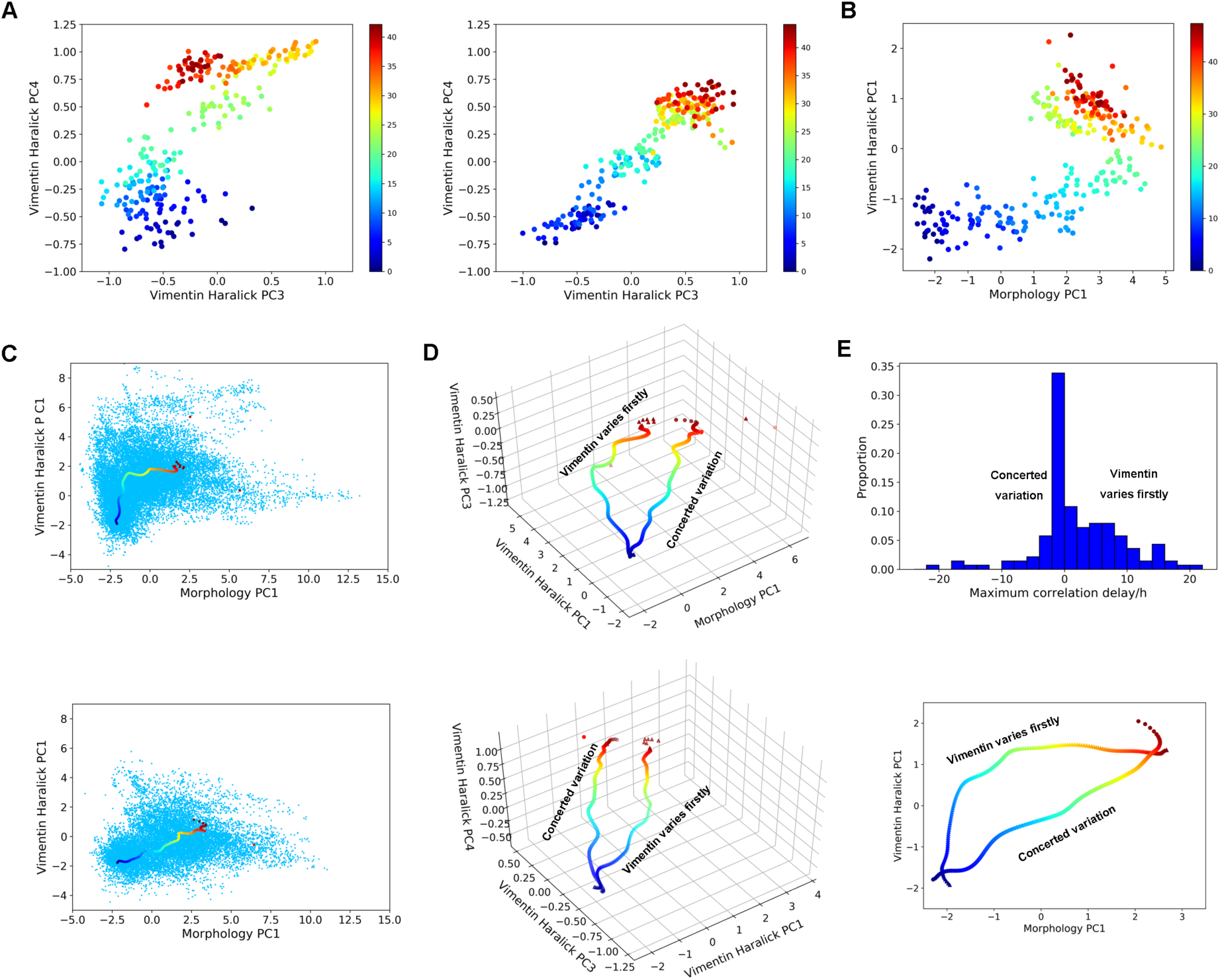
Additional results for EMT path analysis. **(A)** The single EMT trajectory shown in Fig. 6A in the plane of vimentin Haralick PC3 and PC4 (left) and the single EMT trajectory in Fig.6B on the plane of vimentin Haralick PC3 and PC4 (right). Color represents time in hour. **(B)** A single EMT trajectory in which morphology PC1 varies much earlier than vimentin Haralick PC1. Color represents time (unit in hour). **(C)** Scatter plot of all reactive Class I trajectories and the corresponding mean trajectory (top) and scatter plot of all reactive Class I trajectories and the corresponding mean trajectory (bottom). **(D)** Comparison between the mean trajectories of the two classes in the 3D domain of morphology PC1, vimentin Haralick PC1 and PC3 (top), and in the 3D domain of vimentin Haralick PC1, PC3 and PC4 (bottom). **(E)** Distribution of reactive trajectories of time delay with maximum cross correlation between morphology PC1 and vimentin Haralick PC1 (top). Mean trajectories (using the soft-DTW barycenter method) of two groups of trajectories classified by the time delay with maximum cross correlation between morphology PC1 and vimentin Haralick PC1 (bottom).

**Figure S9.**
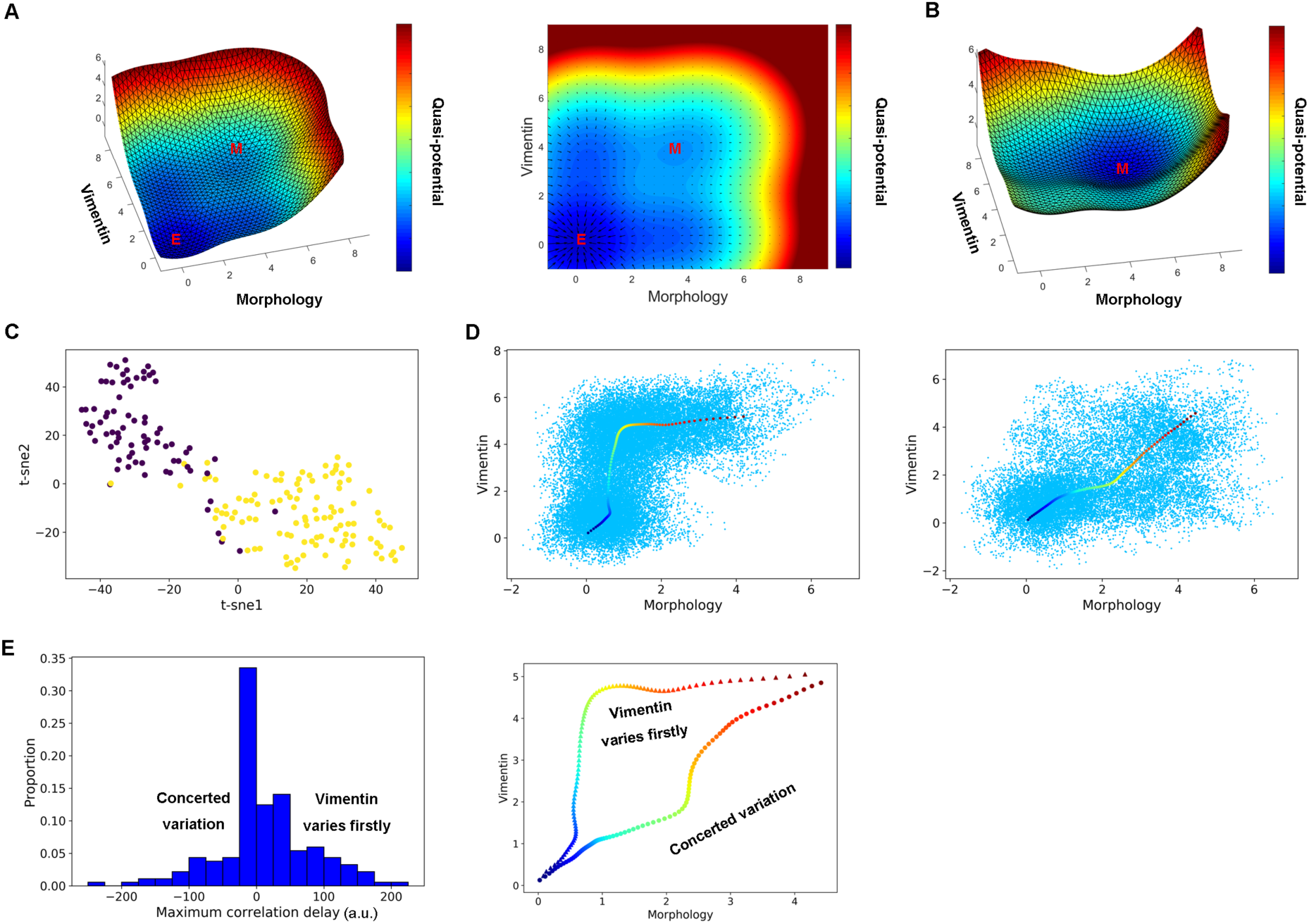
Additional results of simulation of EMT network model. (**A**) Quasi-potential landscape (left) and vector field (right) of EMT network model without TGF-β treatment. (**B**) Quasi-potential landscape of EMT network model with TGF-β treatment. (**C**) Projection of simulated reactive trajectories on 2D t-SNE space by using the DTW distances. Color represents labels of k-means clustering on the DTW distances. (**D**) Scattered plot of all simulated Class I trajectories and the corresponding mean trajectory (left), and scattered plot of all simulated Class II trajectories and the corresponding mean trajectory (right). (**E)** Distribution of simulated trajectories of time delay with maximum cross correlation between morphology and vimentin (left), and corresponding mean trajectories of two groups of trajectories classified by the time delay with maximum cross correlation between morphology and vimentin (using the soft-DTW barycenter method, right).

**Movie S1: A recorded example live cell EMT trajectory**. Each frame is a segmented cell mask cropped from the original recorded vimentin fluorescence image.

**Movie S2: Movie of morphology outlines of the EMT trajectory in Movie S1.** (top) and its trajectory in the plane of morphology PC1 and vimentin Haralick PC1 (bottom).

## References

1. M. A. Nieto, R. Y. Huang, R. A. Jackson, J. P. Thiery, Emt: 2016. Cell 166, 21–45 (2016).

2. K. Takahashi, S. Yamanaka, Induction of Pluripotent Stem Cells from Mouse Embryonic and Adult Fibroblast Cultures by Defined Factors. Cell 126, 663–676 (2006).

3. P. Hanggi, P. Talkner, M. Borkovec, Reaction-rate theory: 50 years after Kramers. Rev. Mod. Phys. 62, 254–341 (1990).

4. W. E. E. Vanden-Eijnden, Transition-path theory and path-finding algorithms for the study of rare events. Annu Rev Phys Chem 61, 391–420 (2010).

5. C. Trapnell, D. Cacchiarelli, J. Grimsby, P. Pokharel, S. Li, M. Morse, N. J. Lennon, K. J. Livak, T. S. Mikkelsen, J. L. Rinn, The dynamics and regulators of cell fate decisions are revealed by pseudotemporal ordering of single cells. Nature biotechnology 32, 381 (2014).

6. R. Kafri, J. Levy, M. B. Ginzberg, S. Oh, G. Lahav, M. W. Kirschner, Dynamics extracted from fixed cells reveal feedback linking cell growth to cell cycle. Nature 494, 480–483 (2013).

7. G. La Manno, R. Soldatov, A. Zeisel, E. Braun, H. Hochgerner, V. Petukhov, K. Lidschreiber, M. E. Kastriti, P. Lönnerberg, A. Furlan, J. Fan, L. E. Borm, Z. Liu, D. van Bruggen, J. Guo, X. He, R. Barker, E. Sundström, G. Castelo-Branco, P. Cramer, I. Adameyko, S. Linnarsson, P. V. Kharchenko, RNA velocity of single cells. Nature 560, 494–498 (2018).

8. C. Weinreb, S. Wolock, B. K. Tusi, M. Socolovsky, A. M. Klein, Fundamental limits on dynamic inference from single-cell snapshots. Proc Natl Acad Sci USA 115, E2467–E2476 (2018).

9. J. Selimkhanov, B. Taylor, J. Yao, A. Pilko, J. Albeck, A. Hoffmann, L. Tsimring, R. Wollman, Accurate information transmission through dynamic biochemical signaling networks. Science 346, 1370–1373 (2014).

10. L. Goentoro, O. Shoval, M. W. Kirschner, U. Alon, The incoherent feedforward loop can provide fold-change detection in gene regulation. Mol Cell 36, 894–899 (2009).

11. S. Gordonov, M. K. Hwang, A. Wells, F. B. Gertler, D. A. Lauffenburger, M. Bathe, Time series modeling of live-cell shape dynamics for image-based phenotypic profiling. Integr Biol (Camb) 8, 73–90 (2016).

12. J. L. McFaline-Figueroa, A. J. Hill, X. Qiu, D. Jackson, J. Shendure, C. Trapnell, A pooled single-cell genetic screen identifies regulatory checkpoints in the continuum of the epithelial-to-mesenchymal transition. Nature Genetics 51, 1389–1398 (2019).

13. L. G. Karacosta, B. Anchang, N. Ignatiadis, S. C. Kimmey, J. A. Benson, J. B. Shrager, R. Tibshirani, S. C. Bendall, S. K. Plevritis, Mapping lung cancer epithelial-mesenchymal transition states and trajectories with single-cell resolution. Nat Commun 10, 5587 (2019).

14. J. Zhang, X. J. Tian, H. Zhang, Y. Teng, R. Li, F. Bai, S. Elankumaran, J. Xing, TGF-β-induced epithelial-to-mesenchymal transition proceeds through stepwise activation of multiple feedback loops. Science signaling 7, ra91 (2014).

15. C. Li, J. Wang, Quantifying Waddington landscapes and paths of non-adiabatic cell fate decisions for differentiation, reprogramming and transdifferentiation. J R Soc Interface 10, 20130787 (2013).

16. P. G. Bolhuis, D. Chandler, C. Dellago, P. L. Geissler, Transition path sampling: Throwing Ropes Over Rough Mountain Passes, in the Dark. Ann. Rev. Phys. Chem. 53, 291–318 (2002).

17. T. F. Cootes, C. J. Taylor, D. H. Cooper, J. Graham, Active shape models-their training and application. Computer vision and image understanding 61, 38–59 (1995).

18. K. Keren, Z. Pincus, G. M. Allen, E. L. Barnhart, G. Marriott, A. Mogilner, J. A. Theriot, Mechanism of shape determination in motile cells. Nature 453, 475–480 (2008).

19. G. J. Stephens, B. Johnson-Kerner, W. Bialek, W. S. Ryu, Dimensionality and dynamics in the behavior of C. elegans. PLoS Comput Biol 4, e1000028 (2008).

20. Z. Pincus, J. Theriot, Comparison of quantitative methods for cell-shape analysis. Journal of microscopy 227, 140–156 (2007).

21. J. Zhang, H. Chen, R. Li, D. A. Taft, G. Yao, F. Bai, J. Xing, Spatial clustering and common regulatory elements correlate with coordinated gene expression. PLOS Comp Biol. 15, e1006786 (2019).

22. J. Maier, B. Traenkle, U. Rothbauer, Visualizing Epithelial–Mesenchymal Transition Using the Chromobody Technology. Cancer research 76, 5592–5596 (2016).

23. L. Zhang, W. Min, Bioorthogonal chemical imaging of metabolic changes during epithelial–mesenchymal transition of cancer cells by stimulated Raman scattering microscopy. Journal of biomedical optics 22, 106010 (2017).

24. N. Costigliola, L. Ding, C. J. Burckhardt, S. J. Han, E. Gutierrez, A. Mota, A. Groisman, T. J. Mitchison, G. Danuser, Vimentin fibers orient traction stress. Proceedings of the National Academy of Sciences 114, 5195–5200 (2017).

25. J. Wang, X. Zhou, P. L. Bradley, S.-F. Chang, N. Perrimon, S. T. Wong, Cellular phenotype recognition for high-content RNA interference genome-wide screening. Journal of Biomolecular Screening 13, 29–39 (2008).

26. J. C. Caicedo, S. Cooper, F. Heigwer, S. Warchal, P. Qiu, C. Molnar, A. S. Vasilevich, J. D. Barry, H. S. Bansal, O. Kraus, Data-analysis strategies for image-based cell profiling. Nature methods 14, 849 (2017).

27. J. C. Caicedo, S. Singh, A. E. Carpenter, Applications in image-based profiling of perturbations. Current Opinion in Biotechnology 39, 134–142 (2016).

28. D. Zhou, O. Bousquet, T. N. Lal, J. Weston, B. Schölkopf, in Advances in neural information processing systems. (2004), pp. 321–328.

29. M. Cuturi, M. Blondel, in *Proceedings of the 34th International Conference on Machine Learning-Volume 70*. (JMLR. org, 2017), pp. 894–903.

30. L. v. d. Maaten, G. Hinton, Visualizing data using t-SNE. Journal of machine learning research 9, 2579–2605 (2008).

31. R. Tavenard, tslearn: A machine learning toolkit dedicated to time-series data, 2017. *URL* https://github.com/rtavenar/tslearn.

32. M. Rhudy, B. Bucci, J. Vipperman, J. Allanach, B. Abraham, in ASME 2009 International Mechanical Engineering Congress and Exposition. (American Society of Mechanical Engineers, 2009), pp. 281–288.

33. Q. Zhong, A. G. Busetto, J. P. Fededa, J. M. Buhmann, D. W. Gerlich, Unsupervised modeling of cell morphology dynamics for time-lapse microscopy. Nat Methods 9, 711–713 (2012).

34. M. Held, M. H. Schmitz, B. Fischer, T. Walter, B. Neumann, M. H. Olma, M. Peter, J. Ellenberg, D. W. Gerlich, CellCognition: time-resolved phenotype annotation in high-throughput live cell imaging. Nat Methods 7, 747–754 (2010).

35. X. Ma, O. Dagliyan, K. M. Hahn, G. Danuser, Profiling cellular morphodynamics by spatiotemporal spectrum decomposition. PLoS computational biology 14, e1006321 (2018).

36. H. Shafqat-Abbasi, J. M. Kowalewski, A. Kiss, X. Gong, P. Hernandez-Varas, U. Berge, M. Jafari-Mamaghani, J. G. Lock, S. Stromblad, An analysis toolbox to explore mesenchymal migration heterogeneity reveals adaptive switching between distinct modes. Elife 5, e11384 (2016).

37. Z. Yin, A. Sadok, H. Sailem, A. McCarthy, X. Xia, F. Li, M. A. Garcia, L. Evans, A. R. Barr, N. Perrimon, C. J. Marshall, S. T. Wong, C. Bakal, A screen for morphological complexity identifies regulators of switch-like transitions between discrete cell shapes. Nat Cell Biol 15, 860–871 (2013).

38. J. E. Sero, H. Z. Sailem, R. C. Ardy, H. Almuttaqi, T. Zhang, C. Bakal, Cell shape and the microenvironment regulate nuclear translocation of NF-B in breast epithelial and tumor cells. Molecular Systems Biology 11, 790–790 (2015).

39. W. Wang, D. A. Taft, Y.-J. Chen, J. Zhang, C. T. Wallace, M. Xu, S. C. Watkins, J. Xing, Learn to segment single cells with deep distance estimator and deep cell detector. Computers in Biology and Medicine 108, 133–141 (2019).

40. K. Jaqaman, D. Loerke, M. Mettlen, H. Kuwata, S. Grinstein, S. L. Schmid, G. Danuser, Robust single-particle tracking in live-cell time-lapse sequences. Nature methods 5, 695 (2008).

41. A. E. Carpenter, T. R. Jones, M. R. Lamprecht, C. Clarke, I. H. Kang, O. Friman, D. A. Guertin, J. H. Chang, R. A. Lindquist, J. Moffat, CellProfiler: image analysis software for identifying and quantifying cell phenotypes. Genome biology 7, R100 (2006).

42. N. Zayed, H. A. Elnemr, Statistical analysis of haralick texture features to discriminate lung abnormalities. Journal of Biomedical Imaging 2015, 12 (2015).

43. M. V. Boland, M. K. Markey, R. F. Murphy, Automated recognition of patterns characteristic of subcellular structures in fluorescence microscopy images. Cytometry: The Journal of the International Society for Analytical Cytology 33, 366–375 (1998).

44. Y. Hu, J. Carmona, R. F. Murphy, in 3rd IEEE International Symposium on Biomedical Imaging: Nano to Macro, 2006. (IEEE, 2006), pp. 1028–1031.

45. L. P. Coelho, Mahotas: Open source software for scriptable computer vision. *arXiv preprint arXiv:1211*.4907, (2012).

46. R. M. Haralick, Statistical and structural approaches to texture. Proceedings of the IEEE 67, 786–804 (1979).

47. C. L. Frick, C. Yarka, H. Nunns, L. Goentoro, Sensing relative signal in the Tgf-β/Smad pathway. Proc Natl Acad Sci U S A 114, E2975–E2982 (2017).

48. R. E. Lee, S. R. Walker, K. Savery, D. A. Frank, S. Gaudet, Fold change of nuclear NF-kappaB determines TNF-induced transcription in single cells. Mol Cell 53, 867–879 (2014).

49. Q. Li, Y. Zheng, X. Xie, Y. Chen, W. Liu, W.-Y. Ma, in Proceedings of the 16th ACM SIGSPATIAL international conference on Advances in geographic information systems. (ACM, 2008), pp. 34.

50. S. Lamouille, R. Derynck, Cell size and invasion in TGF-β–induced epithelial to mesenchymal transition is regulated by activation of the mTOR pathway. The Journal of cell biology 178, 437–451 (2007).

51. L. Buitinck, G. Louppe, M. Blondel, F. Pedregosa, A. Mueller, O. Grisel, V. Niculae, P. Prettenhofer, A. Gramfort, J. Grobler, API design for machine learning software: experiences from the scikit-learn project. *arXiv preprint arXiv:1309*.0238, (2013).

52. E. Jones, T. Oliphant, P. Peterson, SciPy: Open source scientific tools for Python. (2001).

53. S. Chib, E. Greenberg, Understanding the metropolis-hastings algorithm. The american statistician 49, 327–335 (1995).

54. N.-G. Kim, E. Koh, X. Chen, B. M. Gumbiner, E-cadherin mediates contact inhibition of proliferation through Hippo signaling-pathway components. Proceedings of the National Academy of Sciences 108, 11930–11935 (2011).

55. J. T. Morgan, C. J. Murphy, P. Russell, What do mechanotransduction, Hippo, Wnt, and TGFβ have in common? YAP and TAZ as key orchestrating molecules in ocular health and disease. Experimental eye research 115, 1–12 (2013).

56. D.-E. Pefani, D. Pankova, A. G. Abraham, A. M. Grawenda, N. Vlahov, S. Scrace, E. O’Neill, TGF-β targets the hippo pathway scaffold RASSF1A to facilitate YAP/SMAD2 nuclear translocation. Molecular cell 63, 156–166 (2016).

57. C.-Y. Liu, H.-H. Lin, M.-J. Tang, Y.-K. Wang, Vimentin contributes to epithelial-mesenchymal transition cancer cell mechanics by mediating cytoskeletal organization and focal adhesion maturation. Oncotarget 6, 15966 (2015).

58. R. Virtakoivu, A. Mai, E. Mattila, N. De Franceschi, S. Y. Imanishi, G. Corthals, R. Kaukonen, M. Saari, F. Cheng, E. Torvaldson, Vimentin–ERK Signaling Uncouples Slug Gene Regulatory Function. Cancer research 75, 2349–2362 (2015).

59. K. Vuoriluoto, H. Haugen, S. Kiviluoto, J. Mpindi, J. Nevo, C. Gjerdrum, C. Tiron, J. Lorens, J. Ivaska, Vimentin regulates EMT induction by Slug and oncogenic H-Ras and migration by governing Axl expression in breast cancer. Oncogene 30, 1436 (2011).

60. J. Ivaska, Vimentin: Central hub in EMT induction? Small GTPases 2, 1436–1448 (2011).

61. M. G. Mendez, S.-I. Kojima, R. D. Goldman, Vimentin induces changes in cell shape, motility, and adhesion during the epithelial to mesenchymal transition. The FASEB Journal 24, 1838–1851 (2010).

62. J. X. Zhou, M. Aliyu, E. Aurell, S. Huang, Quasi-potential landscape in complex multi-stable systems. Journal of the Royal Society Interface 9, 3539–3553 (2012).

63. J. Xing, Mapping between dissipative and Hamiltonian systems. Journal of Physics A: Mathematical and Theoretical 43, 375003 (2010).

